# Molecular insights into the dual activation of PomZ, a ParA/MinD P-loop ATPase, by the two ATPase activating proteins PomX and PomY

**DOI:** 10.1101/2025.12.01.691545

**Authors:** Philipp Klos, Andrea Harms, Lotte Søgaard-Andersen

## Abstract

Positioning of the division site is precisely regulated. In *Myxococcus xanthus*, the PomX/Y/Z assembly associates with and translocates across the nucleoid in a PomZ ATPase-dependent manner to position the cytokinetic FtsZ-ring at midcell. Unlike other systems incorporating a ParA/MinD ATPase, the PomX/Y/Z system incorporates two ATPase Activating Proteins (AAPs), PomX and PomY, that stimulate PomZ activity. While PomX AAP activity, similarly to other characterized AAPs of ParA/MinD ATPases, resides in a short positively charged N-terminal peptide (PomX^NPEP^), the mechanism of PomY-mediated activation of PomZ remained unknown. We map PomY AAP activity to its C-terminal residues 496-682 (PomY^496-682^) and show that this activity is important for division site positioning and the initiation of cell division. Structural modeling suggests a bipartite PomY^496-682^ architecture, comprising a globular domain and an intrinsically disordered region containing the short α5 helix. However, modeling supported an interaction only between α5 and the PomZ dimer. The α5 binding site partially overlaps with that of the prototype AAP ParB on the ParA dimer but α5 lacks a positively charged residue critical for ParB AAP activity. Modeling supported that PomX^NPEP^ binds the PomZ dimer at a site distinct from that of α5 and which overlaps with that of the prototype AAP MinE on the MinD dimer. These findings reveal a non-canonical AAP mechanism of PomY, while PomX^NPEP^ likely functions analogously to MinE, indicating that PomX and PomY act in concert through distinct but complementary mechanisms to stimulate PomZ ATPase activity, thereby efficiently driving the positioning of the cytokinetic FtsZ-ring.

**Importance:** In bacteria, the regulatory systems that position the division site often incorporate a ParA/MinD ATPase. These ATPases also function in the positioning of numerous other macromolecular complexes in bacteria. Characteristically, these ATPases depend on stimulation of their ATPase activity by a single cognate ATPase activating protein (AAP). However, the *Myxococcus xanthus* cell division regulatory system PomX/Y/Z system stands out by incorporating two AAPs, PomX and PomY, that activate the PomZ ATPase. Similar to other AAPs, PomX uses a short, positively charged N-terminal peptide (PomX^NPEP^) to stimulate PomZ activity. We map PomY AAP activity to a C-terminal region spanning residues 496-682 (PomY^496-682^). Structural modeling using AlphaFold 3 supports that the AAP activity of PomY^496-682^ and PomX^NPEP^ involves two distinct, non-overlapping binding sites at the dimer interface of ATP-bound PomZ. Altogether, these results provide insights into the molecular mechanism of the activation by two AAPs of a ParA/MinD ATPase.

## Introduction

Correct positioning of the cell division site is spatiotemporally precisely regulated to ensure the correct size, shape, and chromosome content of daughter cells. In most bacteria, cell division is initiated by the GTP-dependent polymerization of the tubulin homolog FtsZ at the nascent division site (1, 2). This polymerization results in the formation of a dynamic, ring-like structure, the so-called (Fts)Z-ring, consisting of short treadmilling FtsZ filaments that directly or indirectly recruit the remaining proteins of the division machinery that executes cytokinesis (1, 2). Accordingly, the diverse systems that regulate division site positioning in bacteria spatiotemporally regulate Z-ring formation (3). Several of these systems incorporate a member of the ParA/MinD family of P-loop ATPases that interacts with system-specific components (2, 4, 5). Among the best-studied of these systems, the *Escherichia coli* MinCDE system, the *Bacillus subtilis* MinCDJ/DivIVA system, and the *Caulobacter crescentus* MipZ/ParB system inhibit Z-ring formation at and close to the cell poles, thereby indirectly facilitating Z-ring formation at the nascent division site at midcell (2, 4, 5), while the *Myxococcus xanthus* PomX/Y/Z system directly recruits FtsZ to the nascent division site (6–8).

ParA/MinD ATPases are not only important for the positioning of the division site but also function in chromosome and plasmid segregation as well as in the positioning of diverse macromolecular complexes including carboxysomes, flagella, chemoreceptor arrays and the conjugation machinery (4, 5). The function of ParA/MinD ATPases critically depends on the ATPase cycle during which they alternate between a monomeric state and an ATP-bound dimeric state (9–15). In the symmetric dimer, two ATP molecules are sandwiched between the two subunits (9–14). In the case of MinD homologs, the ATP-bound dimer interacts with the cytoplasmic membrane *via* a C-terminal α-helix (16, 17), while dimeric ParA homologs bind DNA non-specifically *via* a patch of positively charged residues at the dimer interface (9, 14, 18–21). Upon ATP hydrolysis, the dimer dissociates and the ADP-bound monomers are released from the cytoplasmic membrane/DNA and then undergoes spontaneous nucleotide exchange to regenerate the ATP-bound dimer (9, 11, 22, 23).

Although ParA/MinD ATPases contain all residues necessary for ATP hydrolysis, they have a low intrinsic ATPase activity, and they functionally depend on stimulation of their ATPase activity by a cognate ATPase activating protein (AAP) (7, 9, 15, 20, 23–34). While these AAPs are much less conserved than the ParA/MinD ATPases, they share in common that stimulation of their cognate ATPase occurs *via* a short peptide located at or close to the N-terminus and in which one or more positively charged residues are essential for activity (9, 23, 24, 26, 27, 30, 32). Among the best-characterized AAPs, ParB- and MinE-type AAPs function with ParA proteins in chromosome and plasmid segregation and MinD proteins in division site positioning, respectively (4, 5). Interestingly, recent work has revealed that their short, positively charged, N-terminal peptides bind at two distinct sites at the ParA/MinD dimer interface to activate their partner ATPase (26, 35). With the exception of the PomX/Y/Z system (7), these systems incorporate a single AAP. Here, we focus on the activation of the ATPase activity of the DNA-binding PomZ ATPase by the two AAPs PomY and PomX.

The PomX/Y/Z system positions the division site at midcell between two fully segregated chromosomes in the rod-shaped *M. xanthus* cells (7, 8). *In vivo,* PomX forms a single, large meshwork of randomly intertwined self-assembled filaments per cell (6, 7, 27, 36). PomY interacts directly with PomX and binds the PomX meshwork (6, 7, 36). As a result, PomY reaches a high local concentration that is above the critical saturation concentration, C*_sat_*, for phase separation, leading to the formation of a single biomolecular condensate *via* surface-assisted condensation on the PomX meshwork (6, 36). ATP-bound dimeric PomZ binds non-specifically to the nucleoid and is recruited to the MDa-sized PomX/Y assembly by both proteins, thereby associating this assembly to the nucleoid (7, 36). The combined PomX/Y/Z assembly has a unique localization pattern over the cell cycle. Immediately after cell division, it is associated with the nucleoid by PomZ close to the new cell pole; driven by the PomX-and PomY-dependent stimulation of PomZ ATPase activity, the assembly translocates across the nucleoid by biased random motion to midcell, where it undergoes constrained motion (7). At midcell, the PomX/Y/Z assembly recruits FtsZ and stimulates Z-ring formation and, thus, initiation of cell division (6, 7). PomY as well as PomZ interacts with FtsZ (7) and *in vitro*, PomY biomolecular condensates enrich FtsZ and stimulate the GTP-dependent polymerization of FtsZ (6). During cytokinesis, the PomX meshwork undergoes binary fission and the two daughters each acquire a smaller PomX meshwork, while the PomY biomolecular condensate disintegrates (6). Over the following cell cycle, the PomX meshwork accretes additional PomX molecules, and the PomY biomolecular condensate reassembles *de novo* on the PomX meshwork (6).

PomX and PomY separately activate the ATPase activity of DNA-bound PomZ and in combination, they synergistically activate this activity (7). In PomX, a short, positively charged 22-residue N-terminal peptide (PomX^NPEP^) is required and sufficient for AAP function (27). By contrast, the mechanism by which PomY fulfils its AAP function remains unknown. Here, we sought to gain molecular and functional insights into the separate and synergistic activation of PomZ ATPase activity. We show that PomY AAP activity resides in a C-terminal region spanning amino acid residues 496-682 (PomY^496-682^) and is important for cell division site positioning and the initiation of cell division. Moreover, PomY^496-682^ and PomX^NPEP^ are sufficient for the synergistic activation of PomZ ATPase activity. Computational structural modeling using AlphaFold 3 (AF3) supports that the AAP activity of PomY^496-682^ and PomX^NPEP^ involves two distinct and non-overlapping binding sites at the dimer interface of ATP-bound PomZ that overlap with the two distinct binding sites of the positively charged N-terminal peptides of ParB and MinE in their cognate ATPases. Altogether, these results provide insights into the molecular mechanism of the synergistic activation by two AAPs of a ParA/MinD ATPase.

## Results

### Computational analysis of PomY

To explore the mechanism of PomY in the activation of PomZ ATPase activity, we structurally modelled monomeric PomY using AF3 (37). The model confirmed that PomY is a multidomain protein, as previously suggested by sequence-based analyses (6). In this model, PomY contains an α-helix from residue 5 to 106 that is predicted to engage in coiled-coil formation (henceforth, the coiled-coil α-helix) (Fig. 1A). This α-helix is followed by a mostly intrinsically disordered region (IDR) that contains three shorter α-helices (α1-3) ranging in length from 11-15 amino acid residues (Fig. 1A). From residue 221 to 428, PomY contains seven HEAT repeats that form an α-solenoid structure (Fig. 1A-B). The HEAT repeats are followed by a mostly IDR that contains two α-helices (α4-5) with lengths of five and nine residues (Fig. 1A). Finally, the C-terminal residues 624-682 form a globular domain consisting of three α-helices and two β-strands in that order (henceforth, the gD domain for globular domain) (Fig. 1A). Overall, the structural model of PomY is of low confidence based on the low predicted template modelling (pTM) score of 0.41 (Fig. S1A). This low value is caused by the large proportion of IDRs, which have low pLDDT scores Fig. S1A). However, based on pLDDT scores (Fig. S1A), the coiled-coil α-helix, the HEAT repeats and the gD domain are predicted with high confidence (pLDDT >80), α1, α2 and α5 are predicted with relatively good confidence (pLDDT ∼65), while α3 and α4 are predicted with low confidence (pLDDT <50).

**Figure 1.**
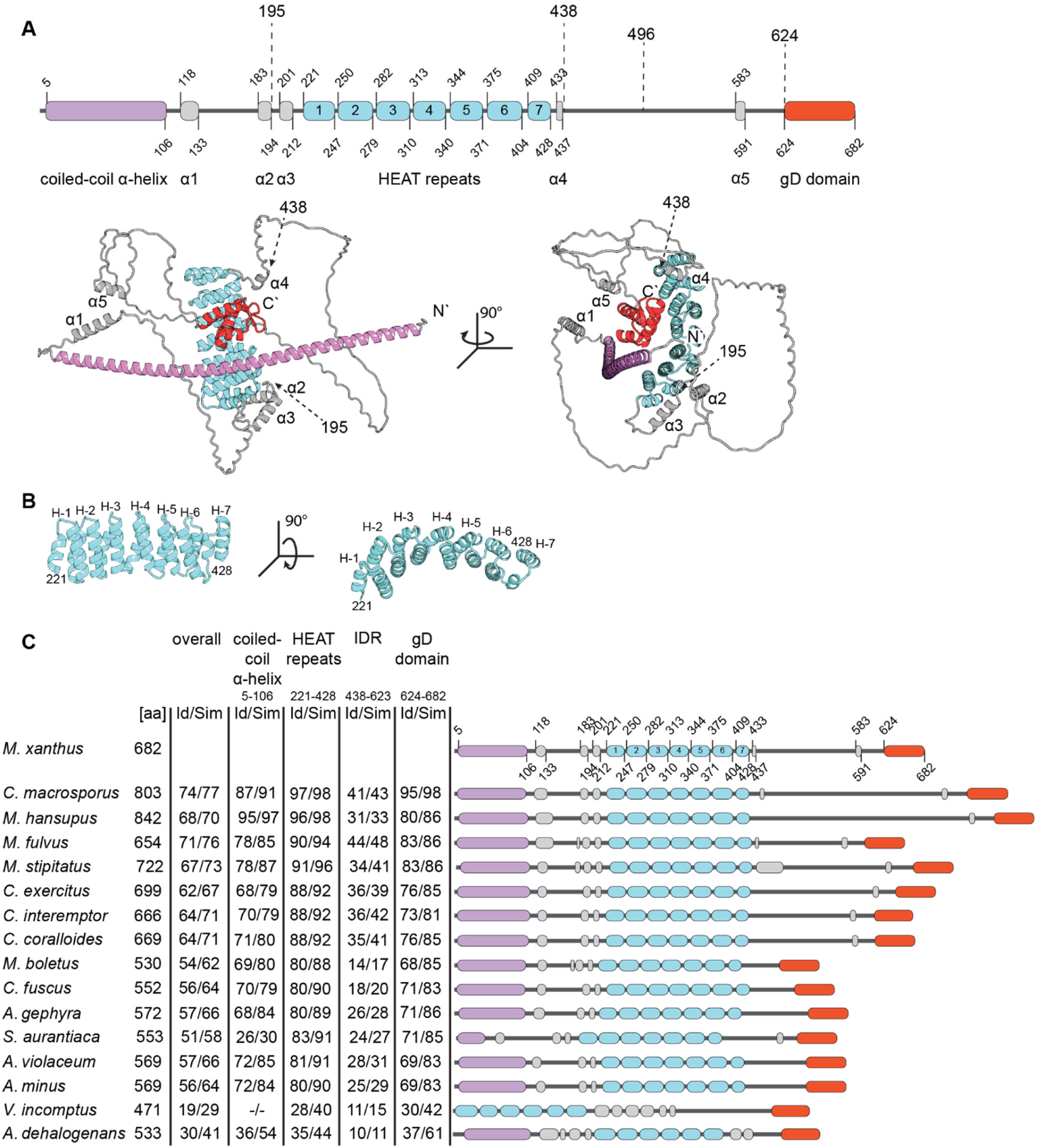
PomY is a multi-domain protein with a conserved domain architecture. (A) PomY domain architecture based on the AF3-based model. Upper panel, schematic of secondary structure elements and the gD domain. Dashed lines indicate the positions to generate truncated PomY variants. Lower panel, AF3 model of PomY with secondary structure elements and the gD domain colored as in the upper panel. Model rank 1 is shown. (B) AF3 model of the seven HEAT-repeats. Individual repeats are labeled H-1-7. (C) The overall PomY architecture is conserved in PomY orthologs. PomY orthologs were identified in a best-best hit reciprocal BlastP analysis from fully sequenced genomes of myxobacteria as indicated on the left. Sequence identity (Id) and similarity (Sim) were calculated based on *M. xanthus* PomY shown in A. Color-code is as in A. Schematics are drawn to scale.

The PomX/Y/Z system is conserved in Myxobacteria (6, 7, 27). Sequence-based and AF3-based analyses demonstrated that the PomY domain architecture is largely conserved among orthologs (Fig. 1C; Fig. S2-S4). In particular, the coiled-coil α-helix, the HEAT repeats and the gD domain are highly conserved (Fig. 1C). By contrast, neither the length nor the sequence of the mostly IDR between the HEAT repeats and the gD domain are highly conserved (Fig. 1C).

We previously demonstrated that PomY self-interacts (6, 7). AF3-based models of PomY modelled an interaction between the coiled-coil α-helix with high confidence in both the dimer and trimer (Fig. S1B-C). These observations support that PomY self-interacts *via* coiled-coil formation; however, it is unclear whether this coiled-coil is composed of two or three α-helices.

### PomY AAP activity resides in the C-terminal mIDR spanning residues 438-682

To examine PomZ ATPase activation by PomY *in vitro*, we purified a His_6_-PomZ variant as previously described (7, 27) (Fig. S5A). Using an NADH-coupled enzymatic assay, we confirmed previous findings (27) that (i) non-specific double stranded herring sperm DNA stimulated His_6_-PomZ ATPase activity, reaching saturation at ∼40µg ml^-1^ (specific (spec.) activity: 6.3±1.0µM ATP hr^-1^ µM^-1^, 4µM His_6_-PomZ) (Fig. S5B) (ii) His_6_-PomZ had no or minimal activity in the absence of DNA at all concentrations tested (spec. activity: 0.3±0.6 ATP hr^-1^ µM^-1^, 4µM His_6_-PomZ) (Fig. S5B-C), and (iii) in the presence of a saturating concentration of DNA (60µg ml^-1^), His_6_-PomZ ATPase activity increased at all concentrations tested without reaching saturation (Fig. S5C).

To investigate PomY AAP activity, we purified two full-length PomY variants. PomY-His_6_, which we previously used to analyze AAP activity (7), and PomY-His_6_-mCherry (from here on PomY-mCh), which we previously used to analyze biomolecular condensate formation by PomY *in vitro* (6) (Fig. S5D). In the absence of DNA, PomY-His_6_ and PomY-mCh only stimulated His_6_-PomZ ATPase activity marginally (spec. activity 1.1±0.8/1.3±0.8 ATP hr^-1^ µM^-1^ at 10µM PomY-His_6_/PomY-mCh, 4µM His_6_-PomZ). However, in the presence of DNA (60µg ml^-1^), PomY-His_6_ and PomY-mCh significantly stimulated His_6_-PomZ ATPase activity, and the stimulation increased with increasing PomY-His_6_/PomY-mCh concentrations, thereby reaching a His_6_-PomZ spec. activity of 10.9±0.9/13.7±3.0 ATP hr^-1^ µM^-1^ at 10µM PomY-His_6_/PomY-mCh and 4µM His_6_-PomZ (Fig. 2A; Fig. S5E). Moreover, His_6_-PomZ ATPase activity did not reach a plateau even at the highest PomY-His_6_ and PomY-mCh concentrations tested. Based on these observations we conclude that DNA and PomY synergistically activate PomZ ATPase activity.

**Figure 2.**
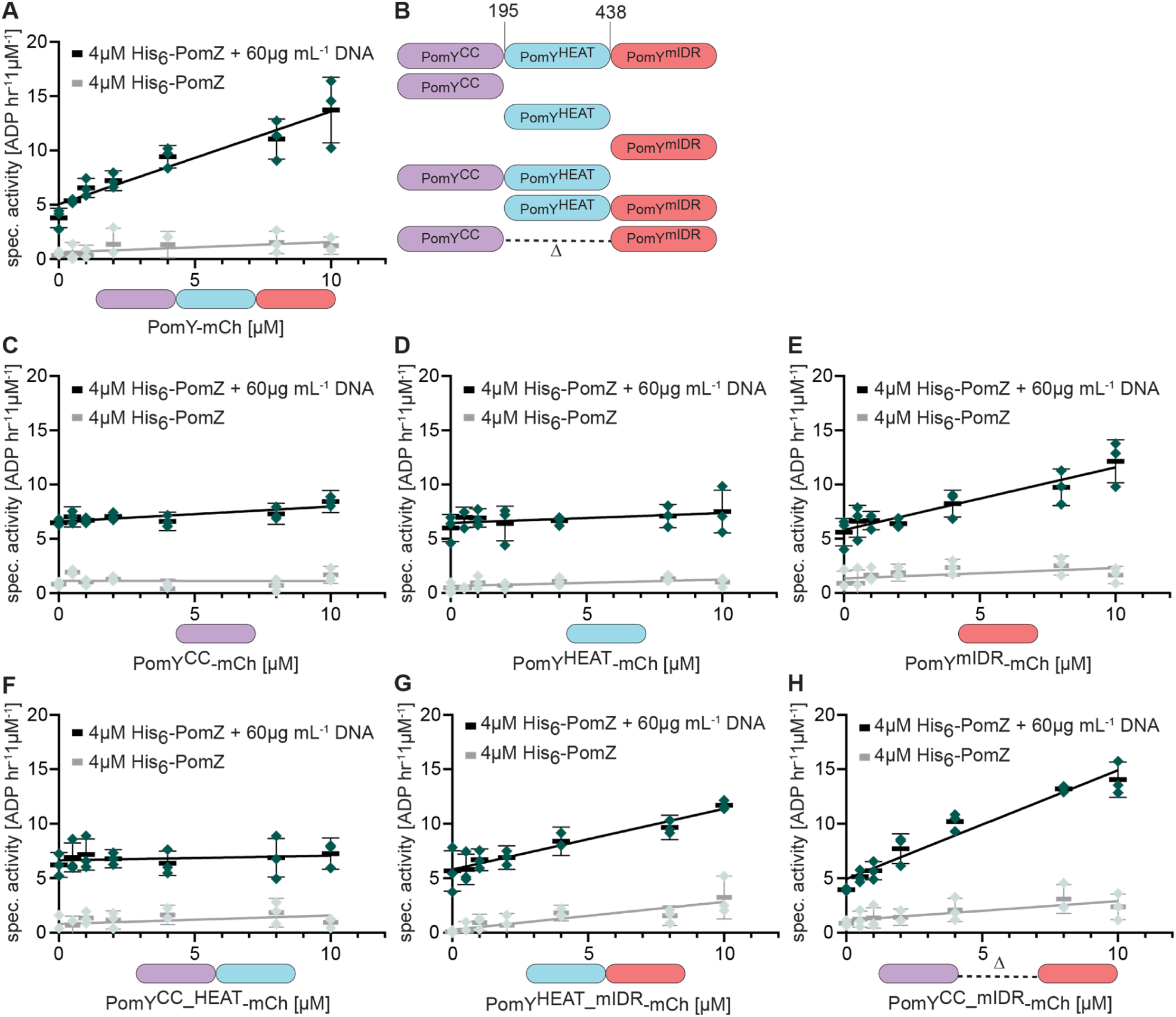
PomY AAP activity resides in the C-terminal mIDR spanning residues 438-682. (A, C-H) Specific His_6_-PomZ ATPase activity. ADP production rate was determined in an NADH-coupled photometric microplate assay in the presence of 1 mM ATP at 32°C. Double stranded herring sperm DNA (DNA) and PomY variants were added as indicated. Green points represent one biological replicate, which was calculated as the mean of three technical replicates. Black bars and error bars, mean of three biological replicates and standard deviation (STDEV). Spontaneous ATP hydrolysis and NADH consumption were accounted for by subtracting the measurements in the absence of His_6_-PomZ. (B) Schematic of truncated PomY variants with truncation points.

To map the part(s) of PomY involved in stimulating PomZ ATPase activity, we divided PomY into three parts (Fig. 1A and 2B). PomY^1-194^ contains the coiled-coil α-helix, α1 and α2 and short IDRs (Fig. 1A) and will be referred to as PomY^CC^ (Fig. 2B). PomY^195-437^ contains the array of seven HEAT repeats, α3 and α4 (Fig. 1A-B) and will be referred to as PomY^HEAT^ (Fig. 2B). PomY^438-682^ contains a long IDR, which is interrupted by the short α5, and the gD domain (Fig. 1A) and will be referred to as PomY^mIDR^, for mostly IDR (Fig. 2B). The truncated PomY variants were purified as mCh-His_6_-tagged proteins (Fig. S5D).

Neither PomY^CC^-mCh nor PomY^HEAT^-mCh measurably stimulated His_6_-PomZ ATPase activity (Fig. 2C-D). By contrast, PomY^mIDR^-mCh stimulated His_6_-PomZ ATPase activity as efficiently as full-length PomY-mCh (spec. activity: 12.1±2.0 ATP hr^-1^ µM^-1^ at 10µM PomY^mIDR^, 4µM His_6_-PomZ) in a DNA-dependent manner (Fig. 2E). Using PomY variants truncated for only one part (Fig. 2B; Fig. S5D), PomY^CC_HEAT^-mCh did not measurably stimulate His_6_-PomZ ATPase activity (Fig. 2F), while PomY^HEAT_mIDR^-mCh and PomY^CC_mIDR^-mCh stimulated His_6_-PomZ ATPase activity in a DNA-dependent manner as efficiently as full-length PomY-mCh and PomY^mIDR^-mCh (spec. activity PomY^HEAT_mIDR^-mCh/PomY^CC_mIDR^-mCh: 11.7±0.7/14.0±1.6 ATP hr^-1^ µM^-1^ at 10µM PomY-mCh variant, 4µM His_6_-PomZ) (Fig. 2G-H). These findings demonstrate that the C-terminal region spanning residues 438-682 is required and sufficient for stimulating PomZ ATPase activity. From here on, we refer to this region as the PomY^mIDR^ region.

### PomZ interacts with PomY *via* the C-terminal PomY^mIDR^ region

The ATPase assay demonstrates that the PomY^mIDR^ region interacts with PomZ. To determine whether the remaining two parts of PomY interact with PomZ, we analyzed the PomY/PomZ interaction using the yeast-2-hybrid (Y2H) system, as previously described (7). Y2H assays detect protein-protein interactions by reconstituting a functional GAL4 transcription factor, resulting in the expression of the HIS3 reporter gene and growth on histidine deficient medium. Interaction stringency can be increased by adding 3-amino-1,2,4-triazole (3-AT), a competitive inhibitor of the HIS3 gene product.

Using the same truncation points as in the PomY variants used in the ATPase assay (Fig. 2B), we observed at low stringency an interaction between PomZ and PomY^mIDR^ in agreement with the ATPase assay (Fig 3A). Because the strains containing Gal4BD-PomY or Gal4BD-PomY^CC_HEAT^ together with the pGal4AD control plasmid grew at low stringency (Fig. 3A), we increased stringency by adding10mM 3-AT, thereby eliminating this growth (Fig. 3A). At high stringency, we observed PomZ/PomY and PomZ/PomY^mIDR^ interactions but not a PomZ/PomY^CC_HEAT^ interaction. These observations suggest that PomZ only interacts with PomY *via* the C-terminal PomY^mIDR^ region.

**Figure 3.**
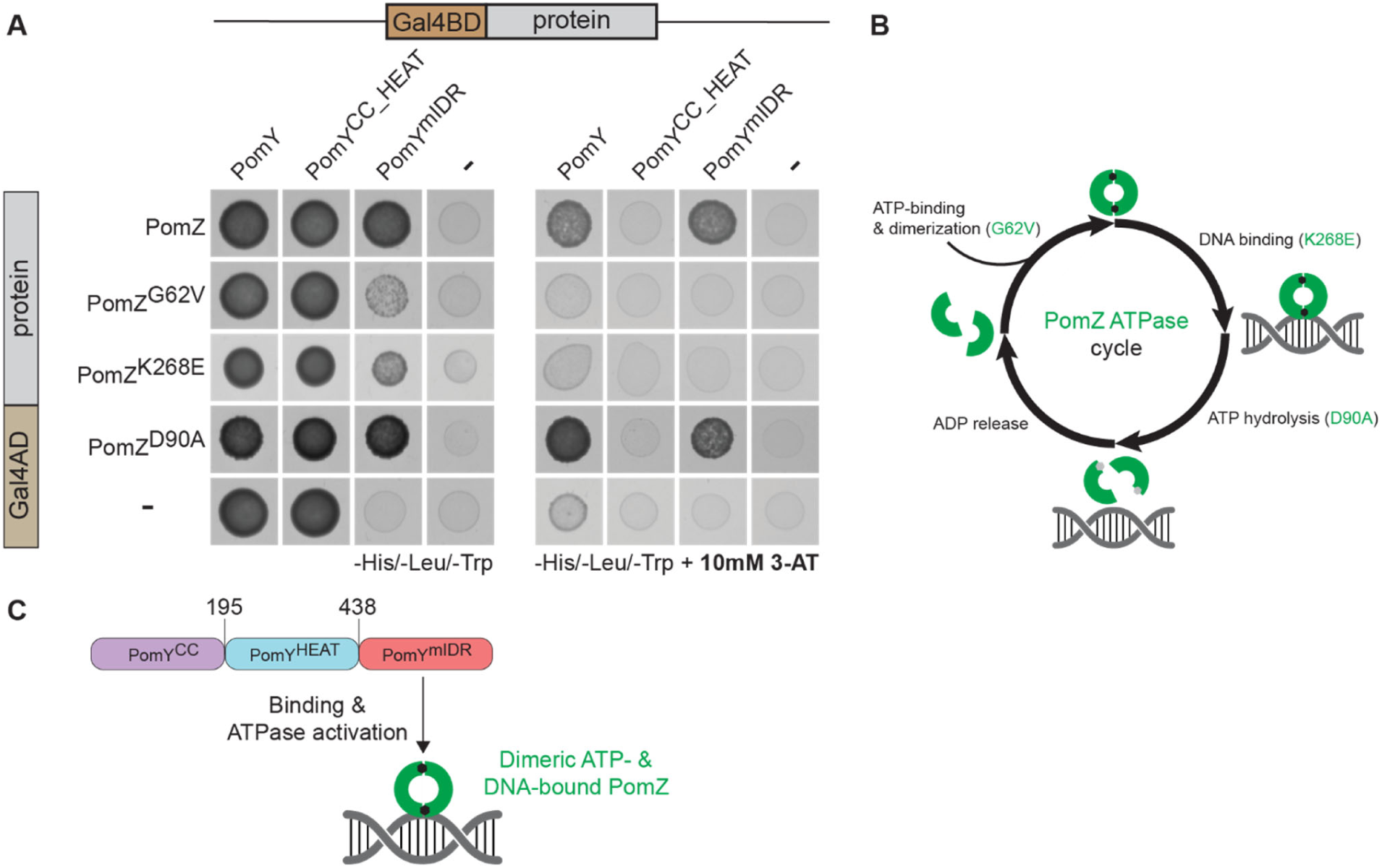
PomY interacts with the PomZ dimer *via* the C-terminal mIDR spanning residues 438-682. (A) Yeast two-hybrid analysis of interaction between PomY and PomZ. Yeast strain AH109 expressing the indicated variants of PomY and PomZ fused to Gal4BD and Gal4AD was analyzed for growth on selective medium. 3-AT was included in plates to increase stringency. Controls were AH109 with a bait plasmid expressing a Gal4AD or Gal4BD fusion and a plasmid expressing only Gal4BD or Gal4AD. Three independent clones were analyzed per strain. (B) ATPase cycle of PomZ. Green semicircles indicate PomZ monomers. PomZ dimerizes upon ATP binding and PomZ^G62V^ is reduced in dimerization; dimeric ATP-bound PomZ binds non-specifically to DNA and PomZ^K268E^ is reduced in DNA binding; dimeric-, DNA-bound PomZ hydrolyzes ATP, resulting in dissociation of the dimer and release from DNA and PomZ^D90A^ is reduced in ATP hydrolysis. (C) Schematic of the interaction between PomY^mIDR^ and dimeric, ATP- and DNA-bound PomZ.

### The ATP-bound dimeric and DNA-binding form of PomZ interacts with the C-terminal terminal PomY^mIDR^ region

To analyze which form of PomZ interacts with PomY, we used a set of previously characterized PomZ variants (7) in which PomZ^G62V^ is reduced in dimerization, PomZ^K268E^ in DNA binding, and PomZ^D90A^ in ATP hydrolysis and is, therefore, locked in the dimeric ATP-bound and DNA-binding form (Fig. 3B). In the Y2H analyses at low stringency, we observed strong interactions of PomZ and PomZ^D90A^ with PomY^mIDR^ and weak interactions of PomZ^G62V^ and PomZ^K268E^ with PomY^mIDR^ (Fig. 3A). At increased stringency in the presence of 10mM 3-AT, PomZ as well as PomZ^D90A^ interacted with PomY and PomY^mIDR^ but not with PomY^CC_HEAT^ (Fig. 3A), and neither PomZ^G62V^ nor PomZ^K268E^ interacted with PomY, PomY^CC_HEAT^ and PomY^mIDR^ (Fig. 3A). These observations strongly support that the PomY^mIDR^ region specifically interacts with the dimeric ATP-bound and DNA-binding form of PomZ (Fig. 3C).

### PomY AAP activity resides in the C-terminal PomY^496-682^ region

Having established that the PomY^mIDR^ region is required and sufficient for the interaction with dimeric PomZ and AAP activity, we asked which subregion(s) are involved in AAP activity. Based on sequence analyses, we divided the PomY^mIDR^ region into three subregions (Fig. 1A; 4A-B). The subregion spanning 438-495 is an IDR and relatively hydrophobic; the subregion spanning 496-623 is also mostly an IDR but includes α5, relatively hydrophilic, and rich in positively charged residues; and, the subregion spanning 624-682 contains the gD domain. To address the function of these three subregions in AAP activity, we aimed to purify full-length PomY mCh-His_6_-tagged variants each lacking one of these subregions (Fig. S6A). Because we were unable to overexpress a protein lacking residues 438-495, we purified a protein lacking residues 438-488 (PomY^Δ438-488^) (Fig. S6A). At 10µM, PomY^Δ438-488^-mCh stimulated His_6_-PomZ ATPase activity as efficiently as full-length PomY in a DNA-dependent manner (Spec. activity: 15.3±1.4 ATP hr^-1^ µM^-1^ at 10µM PomY^Δ438-488^-mCh, 4µM His6-PomZ) (Fig. 4C). By contrast, neither PomY^Δ496-623^-mCh nor PomY^Δ624-682^-mCh significantly stimulated His_6_-PomZ ATPase activity (Fig. 4D-E). Thus, the region spanning residues 496-682 is required for PomY AAP activity. From here on, we refer to this region as the PomY^496-682^ region.

**Figure 4.**
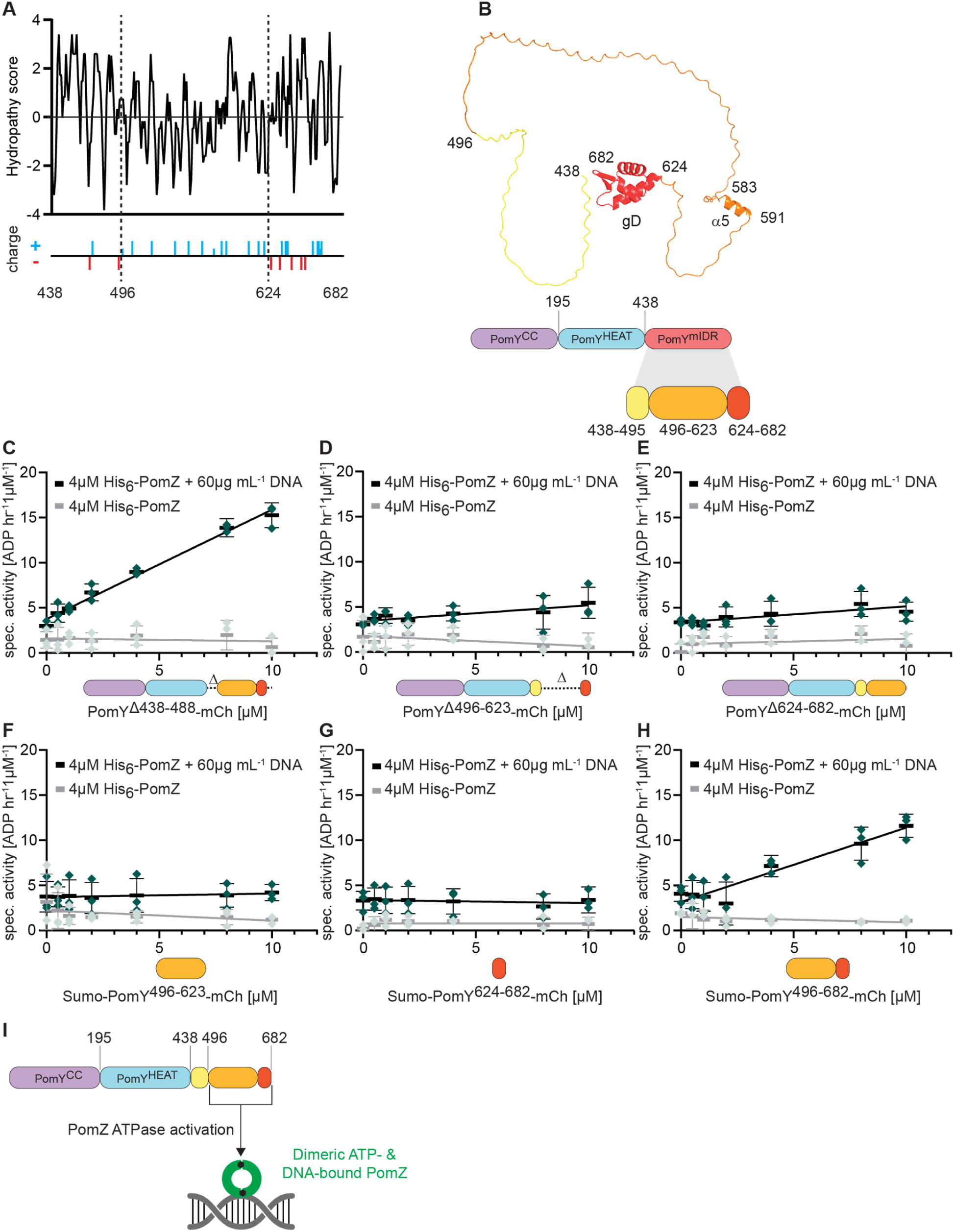
PomY AAP activity resides in the region spanning residues 496-682 in the mIDR. (A) *In silico* characterization of PomY^mIDR^. Upper panel, hydrophobicity analysis of the PomY^mIDR^ sequence. The analysis was performed by the ProtScale web-tool (66) with the parameters of Kyte and Doolittle and a window size of three amino acids (62). Lower panel, positively (blue) and negatively (red) charged amino acid residues. (B) AF3 model and schematic of PomY^mIDR^. Upper panel, AF3 model with subregions colored as in the lower panel. Lower panel, schematic of PomY^mIDR^ with the subregions analyzed in the ATPase assay. (C-H) His_6_-PomZ ATPase activity. DNA and PomY variants were added as indicated. Experiments were performed as in Fig. 2A. (I) Schematic of the interaction between PomY^496-682^ in the mIDR of PomY and dimeric ATP-and DNA-bound PomZ.

To examine whether the PomY^496-682^ region is sufficient for AAP function, we sought to overexpress and purify mCh-His_6_-tagged variants of PomY^496-623^, PomY^624-682^ and PomY^496-682^. Because we were unable to overexpress all these variants, we purified variants with an N-terminal His_6_-Sumo-tag (Sumo) and a C-terminal mCh-tag (Fig. S6B). As a control, we observed that a Sumo-PomY^mIDR^-mCh variant stimulated His_6_-PomZ ATPase activity (spec. activity: 11.0±1.7 ATP hr^-1^ µM^-1^ at 10µM Sumo-PomY^mIDR^-mCh, 4 µM His_6_-PomZ) in a DNA-dependent manner (Fig. S6C) as efficiently as full-length PomY-mCh and PomY^mIDR^-mCh (Fig. 2A, E). Neither Sumo-PomY^496-623^-mCh nor Sumo-PomY^624-682^-mCh stimulated His_6_-PomZ ATPase activity (Fig. 4F-G). However, Sumo-PomY^496-682^-mCh stimulated His_6_-PomZ ATPase activity (spec. activity: 11.6±1.3 ATP hr^-1^ µM^-1^ at 10µM Sumo-PomY^496-682^-mCh, 4 µM PomZ-His_6_) in a DNA-dependent manner (Fig. 4H), similar to the Sumo-PomY^mIDR^-mCh variant (Fig. S6C).

These observations demonstrate that the PomY^496-682^ region is required and sufficient for PomY AAP activity (Fig. 4I).

### Reconstitution of the synergistic activation of PomZ ATPase activity by minimal parts of PomY and PomX

PomX and PomY stimulate PomZ ATPase activity synergistically in the presence of DNA (7). To determine whether PomX^NPEP^, i.e. the positively charged peptide that comprises the N-terminal 22 residues of PomX and is required and sufficient for AAP function (27), and PomY^496-682^ recapitulate this synergistic activation, we incubated 2µM of PomX^NPEP^ and/or PomY^624-682^-mCh with 4µM His_6_-PomZ and DNA. While His_6_-PomZ alone had a spec. activity of 2.9±1.3 ATP hr^-1^ µM^-1^, 2µM PomX^NPEP^ and 2µM PomY^624-682^-mCh stimulated ATPase activity to 6.1±1.5 and 5.1±2.0 ATP hr^-1^ µM^-1^, respectively (Fig. 5). Importantly, in the presence of both 2µM PomX^NPEP^ and 2µM PomY^624-682^-mCh, PomZ had a specific activity of 17.1±1.4 ATP hr^-1^ µM^-1^. Thus, PomX^NPEP^ and PomY^496-682^ are sufficient to activate PomZ ATPase activity synergistically.

**Figure 5.**
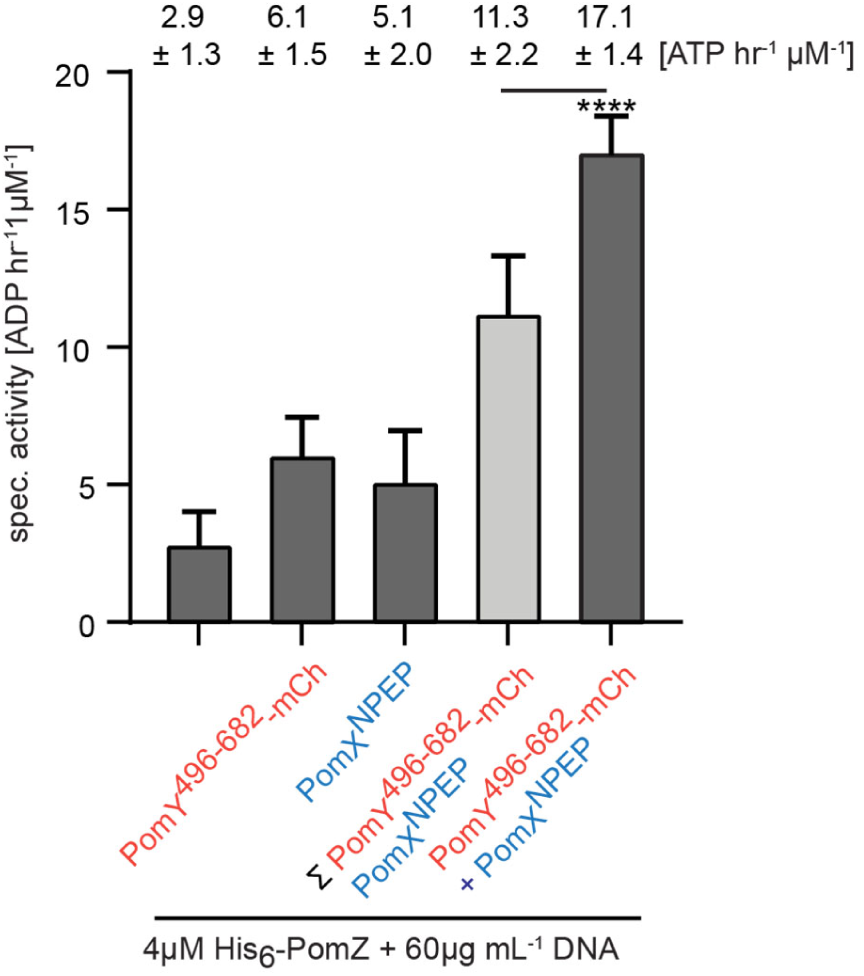
Reconstitution of the synergistic activation of PomZ ATPase activity by minimal parts of PomY and PomX. His_6_-PomZ ATPase activity. Proteins and DNA were added as indicated. PomY^496-682^-mCh and PomX^NPEP^ were added to final concentrations of 2µM. Experiments were performed as in Fig. 2A. Numbers indicate mean spec. activity±STDEV based on three biological replicates, each calculated from three technical replicates. Light grey bar, sum of His_6_-PomZ specific activity in the presence of either PomY^496-682^-mCh or PomX^NPEP^. ****, P <0.0001, unpaired t-test.

### The PomY^496-682^ region is important for division site positioning, initiation of division, and recruitment of PomZ to the PomX/Y assembly

To probe the importance of the PomY^496-682^ region *in vivo*, we fused a PomY variant lacking this region to mCh (PomY^Δ496-682^-mCh), and expressed it as well as wild-type (WT) PomY-mCh ectopically from the native promoter from plasmids integrated in single copies at the Mx8 *attB* site in the Δ*pomY* mutant. The PomY variants accumulated at slightly lower levels than PomY expressed from the native locus (Fig. S7A). WT cells had a cell length of 7.7±0.3 µm (mean±STDEV), while Δ*pomY* cells were filamentous with a length of 14.7±1.5µm and generated minicells (Fig. 6A). PomY-mCh complemented the division defect of the Δ*pomY* mutant in agreement with previous observations (6, 7), while PomY^Δ496-682^-mCh only partially complemented this defect with a length of 9.0±0.7µm and generated minicells (Fig. 6A). Moreover, cells containing PomY^Δ496-682^-mCh had a significantly reduced constriction frequency (Fig. 6B), and while 93% and 89% of the constrictions in WT and the Δ*pomY*/*pomY*-mCh strain occurred at midcell, only 68% of the few constrictions in the Δ*pomY*/*pomY*^Δ496-682^-mCh strain occurred at midcell (Fig. 6C). These observations demonstrate that the PomY^496-682^ region is important for division site positioning at midcell and stimulation of the initiation of cell division. Of note, the results obtained for the PomY^Δ496-682^-mCh variant are similar to those obtained for a PomY variant lacking the entire mIDR (residues 438-682) (PomY^ΔmIDR^-mCh) (6), underscoring that the important functional parts in the PomY^mIDR^ region span residues 496-682.

**Figure 6.**
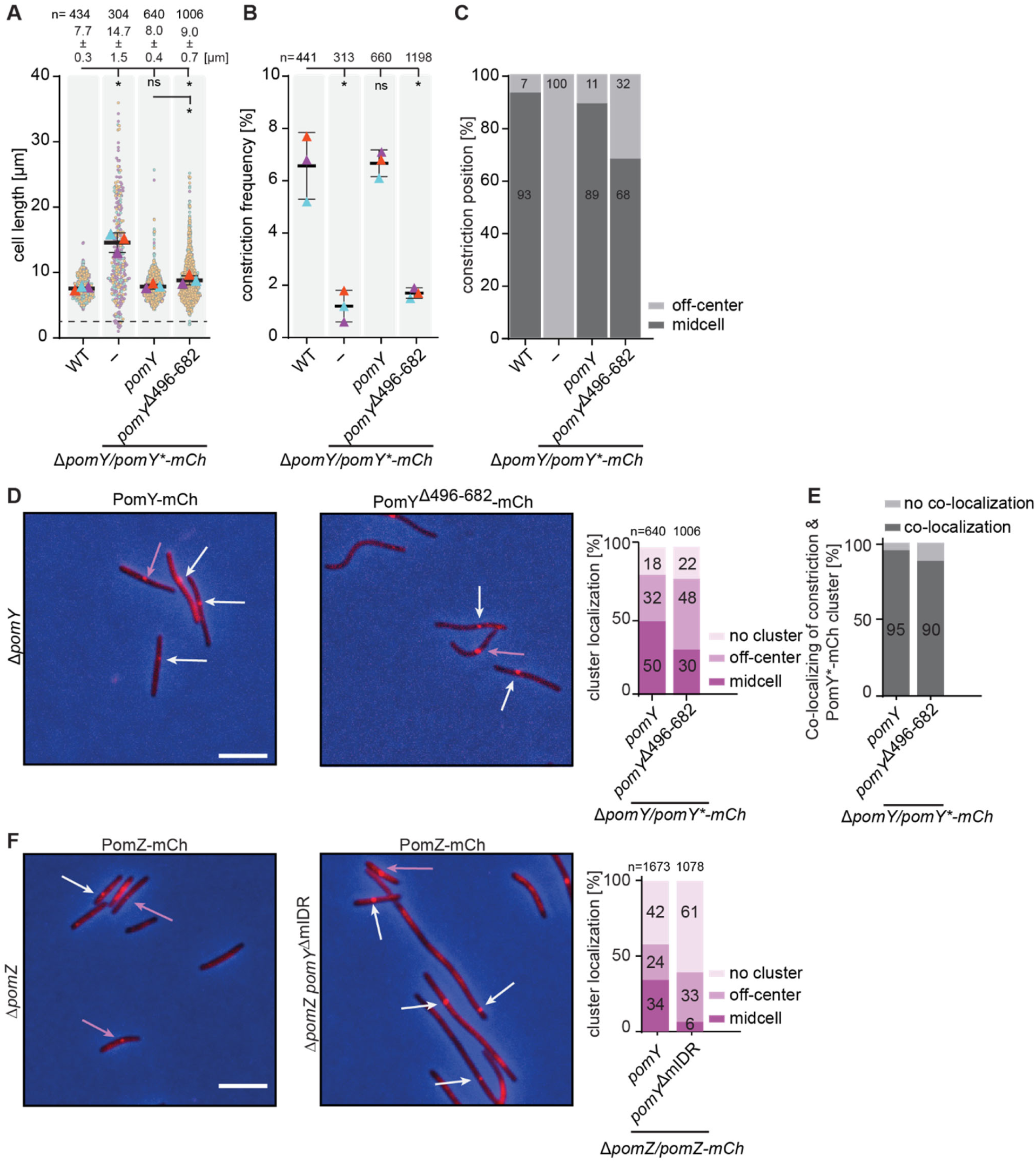
PomY^496-682^ is important for division site positioning at midcell, initiation of cell division, and recruitment of PomZ to the PomX/Y assembly. (A) Cell length distribution of cells of the indicated genotypes. Each strain was imaged in three biological replicates with individual cells and means (triangles) colored according to the replicate. Cells below the stippled line are minicells. Numbers above as well as black bold bars and error bars, mean cell length± STDEV calculated based on the means from the three biological replicates. n, total number of cells analyzed. *, P <0.005, ns, not significant, two-way ANOVA test with multiple comparisons. (B) Constriction frequency of cells of the indicated genotypes. Each strain was imaged in three biological replicates with individual means (triangles) colored according to the replicate. Black bold bars and error bars, mean cell length± STDEV calculated based on the means from the three biological replicates. n, total number of cells analyzed. *, P <0.005, ns, not significant, two-way ANOVA test with multiple comparisons. (C) Constriction position in cells of the indicated genotypes. The same cell as in B were analyzed. Midcell is defined as 50±5% of the cell length. (D, F) Fluorescence microscopy of cells of the indicated genotypes. Left panels, phase-contrast and fluorescence images of representative cells were overlayed; arrows indicate clusters localized at midcell (purple) and off-center (white); scale bar, 5 µm; right diagram, quantification of cluster localization. (E) Cluster/constriction co-localization in cells of the indicated genotypes. The same cells as in B were analyzed.

Next, we determined the localization of PomY^Δ496-682^-mCh by fluorescence microscopy. In agreement with previous observations, PomY-mCh localized in a single cluster at midcell in 50% and in an off-center position in 32% of cells, while 18% of cells did not have a cluster (7). By contrast, PomY^Δ496-682^-mCh localized in a cluster at midcell in 30% of cells, in an off-center position in 48% of cells, and 22% had no cluster (Fig. 6D). Moreover, in the presence of PomY-mCh, 95% of constrictions co-localized with the PomY-mCh cluster, while only 90% of the few constrictions co-localized with the with PomY^Δ496-682^-mCh cluster (Fig. 6E). We conclude that the PomY^496-682^ region is not important for PomY cluster formation, but for cluster localization to midcell.

Next, we determined whether PomZ recruitment to the PomX/Y cluster was reduced in the absence of the PomY^496-682^ region. Because the results for PomY^Δ496-682^-mCh were similar to those obtained for PomY^ΔmIDR^-mCh, we determined the localization of an active PomZ-mCh fusion (7, 8) in the presence of PomY^ΔmIDR^. The truncated PomY^ΔmIDR^ protein accumulated at a level similar to that of full-length PomY (Fig. S7B), while PomZ-mCh abundance was slightly elevated in comparison to the native protein (Fig. S7C). While 58% of WT cells had a visible PomZ-mCh cluster, only 39% of cells had a cluster in the presence of PomY^ΔmIDR^ (Fig. 6F). Moreover, as for PomY^Δ496-682^-mCh, PomZ-mCh cluster localization to midcell was reduced, i.e. 34% of clusters in WT and only 6% of clusters in the presence of PomY^ΔmIDR^ were at midcell, while the frequency of clusters in an off-center position was increased from 24% in WT to 33% in the presence of PomY^ΔmIDR^ (Fig. 6F). Based on these findings and the observation that PomZ interacts with the PomY^496-682^ region *in vitro*, we conclude that this region is important for recruitment of PomZ to the PomX/Y assembly *in vivo*.

### Analysis of the PomY region spanning residues 496-682

Having found that the PomY^496-682^ region contains the PomY AAP activity and is important for PomY function *in vivo*, we performed a detailed computational analysis of this region. The PomY^496-682^ region is mostly an IDR but also contains α5 (Fig. 4B). Sequence-based searches of the region from 624-682 did not identify proteins other than PomY orthologs with the gD domain. A Foldseek search (38) in the PDB database, identified a winged helix-turn-helix (wHTH) domain in the Z-DNA binding Z-α domain of ORF112 from cyprinid herpesvirus 3 (39), an O-methyltransferase from *Burkholderia thailandensis,* the DNA binding domains of the transcriptional regulators MarR of *Listeria monocytogenes* and MepR of *Staphylococcus aureus* (40), and the activation domain of the transcriptional regulator MotA from bacteriophage T4 (41) as the top five structural homologs of the gD domain (Fig. S8A-C).

A multiple sequence alignment revealed that the gD domain lacks the residues involved in Z-DNA binding by characterized Z-α-domains (Fig. S8D). To determine whether the gD domain binds DNA, we used a biolayer interferometry (BLI) assay in which a 226 bp double-biotinylated DNA fragment was immobilized on a streptavidin coated sensor and then probed with PomY^624-682^-mCh, which only contains the gD domain, and His_6_-PomZ as a positive control. We observed concentration-dependent binding by His_6_-PomZ with an apparent equilibrium dissociation constant (K_D_) of 24.3±4.6µM, but not binding by PomY^624-682^-mCh (Fig. S8E). Based on these analyses, we suggest that the gD domain binds neither Z-DNA nor DNA and may engage in protein-protein interaction(s).

### Structural modeling of the PomZ/PomY^496-682^ and the PomZ/PomX^NPEP^ interactions

PomY^496-682^ and PomX^NPEP^ interact with the ATP-bound PomZ dimer and separately as well as synergistically activate PomZ ATPase activity (this study, (27)). To understand how they interact with the PomZ dimer separately and in combination, we performed structural modeling in three steps using AF3. First, we modelled the ATP-bound PomZ dimer (Fig. 7A; Fig S9A). In the high confidence model, each PomZ protomer in the symmetric dimer binds one ATP molecule that is sandwiched between the two protomers. Using Foldseek, we identified the ATP-bound *Helicobacter pylori* ParA (ParA*_Hp_*) dimer bound to DNA (14) as the closest structural homolog (RMSD 1.605 Å over 429 C_α_ atoms) (Fig. S9B). In the ParA*_Hp_* dimer, residues from both protomers form a positively charged, surface-exposed patch that binds DNA (14) (Fig. S9C). A similar patch is present in the PomZ dimer and includes Lys268, which is important for DNA binding by PomZ (7) (Fig. S9D; see also Fig. 3B). Consistently, a high confidence AF3 model of the ATP-bound PomZ dimer and a 25bp DNA fragment showed this positively charged patch in close contact with DNA (Fig. S9E). The PomZ dimers modelled in the absence and presence of DNA aligned almost perfectly (RMSD 0.669 Å over 615 C_α_ atoms) (Fig. S9F). From here on, we used the PomZ dimer model without DNA.

**Fig. 7.**
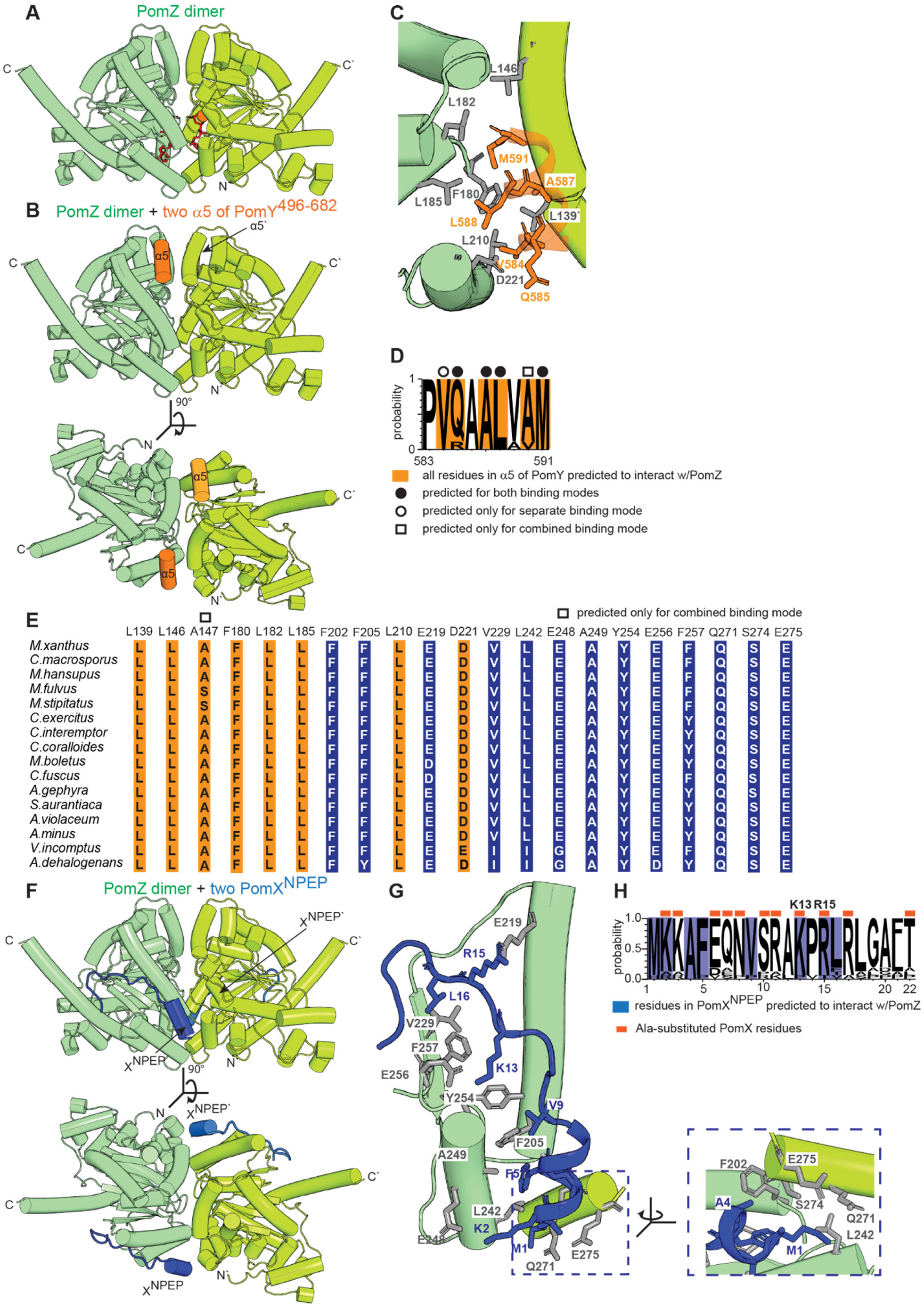
Structural modeling of the PomZ dimer, the PomZ/PomY^496-682^ interaction and the PomZ/PomX^NPEP^ interaction. (A) AF3-based structural model of PomZ dimer. Individual protomers are colored in shades of green. Two ATP molecules are shown in red in stick representation and Mg^2+^ ions in orange. Model rank 1 is shown. (B) AF3-based structural model of PomZ dimer (green shades) with two copies of α5 of PomY^496-682^ (orange). The model was generated with PomY^496-682^, but only the high confidence α5 interactions to the PomZ dimer are included. PomZ was modeled with ATP and Mg^2+^ but the ATP molecules and Mg^2+^ are not included for simplicity. Model rank 1 is shown. (C) Zoom of the predicted interaction of the PomZ dimer and α5 of PomY^496-682^ from the model in B. PomZ residues (grey) and α5 residues (orange) predicted to be involved in the interaction are shown in stick representation. (D) Weblogo consensus sequence of α5 in PomY^496-682^. The weblogo was generated based on the nine PomY orthologs that are predicted to contain an α5 (Fig. 1C). Residues (on orange) predicted to interact with PomZ are based on the models in Fig. 7B-C and Fig. 8A and C; these residues are further marked depending on whether they are predicted to interact with PomZ when PomY^496-682^ is modelled separately with PomZ or PomY^496-682^ is modelled in combination with PomX^NPEP^. (E) Conservation of PomZ residues predicted to interact with α5 of PomY^496-682^ (orange) and PomX^NPEP^ (blue). The indicated conservation is based on the multiple sequence alignment in Fig. S11. (F) AF3-based structural model of PomZ dimer (green shades) with two molecules of PomX^NPEP^ (blue). PomZ was modeled with ATP and Mg^2+^ but the ATP molecules and Mg^2+^ are not included for simplicity. Model rank 1 is shown. (G) Zoom of the predicted interaction of the PomZ dimer and PomX^NPEP^ from the model in F. PomZ residues (grey) and PomX^NPEP^ residues (blue) predicted to be involved in the interaction are shown in stick representation. The blue dashed line indicates the zoomed region in the right panel. (H) Weblogo consensus sequence of PomX^NPEP^. The weblogo was generated based on PomX orthologs from the myxobacteria listed in Fig. 1C. Residues predicted to interact with PomZ are based on the model in F and G as well as Fig. 8A and D and are on blue.

In the second step, we modeled the ATP-bound PomZ dimer with two molecules of either PomY^496-682^ or PomX^NPEP^. In the PomZ/PomY^496-682^ model, based on pLDDT scores, only the PomZ dimer, the two α5’s, and the two gD domains were modelled with high confidence (Fig. S10A). Based on pAE scores, both α5s were modeled with high confidence to interact with the PomZ dimer (Fig. S10A), while the gD domain was neither predicted to interact with the PomZ dimer nor with the remaining parts of PomY^496-682^ (Fig. S10A). Therefore, we focused on the PomZ/α5 interaction. In the model, the two α5s bind the PomZ dimer symmetrically in a mainly hydrophobic cleft between the two protomers opposite the DNA-binding patch (Fig. 7B). Specifically, four hydrophobic residues (V584, A587, L588, M591) and one hydrophilic (Q585) are predicted to face towards the PomZ dimer surface (Fig. 7B-C) and interact with a pocket composed of hydrophobic residues from both PomZ protomers (L139’, L146’, F180, L182, L185, L210) and the charged D221 (Fig. 7C). Remarkably, all nine residues of α5, including the five residues predicted to interact with PomZ, are highly conserved in PomY orthologs that contain an α5 (Fig. 7D), despite the low overall sequence conservation of the region from residues 438-623 (Fig. 1C). Similarly, the PomZ residues in contact distance to α5 are highly conserved in PomZ orthologs but not in ParA and MinD orthologs (Fig. 7E; Fig. S11), thus lending further support to the structural model.

In the PomZ/PomX^NPEP^ model, the PomZ dimer and the two PomX^NPEP^ molecules were modelled with high confidence based on pLDDT scores. Residues 2-8 of PomX^NPEP^ were predicted to be α-helical and the rest unstructured (Fig. 7F). Based on pAE scores (Fig. S10B), both PomX^NPEP^ molecules with high confidence interact symmetrically with the PomZ dimer in a cleft between the two protomers, but at a different position than α5 of PomY (Fig. 7F; cf. Fig. 7B). Specifically, K2, A4 and F5 in the α-helical part and M1, V9, K13, R15, and L16 in the unstructured parts face towards the PomZ dimer surface (Fig. 7G), and are predicted to interact with a pocket composed of residues from both PomZ protomers (F202, F205, E219, V229, L242, E248, A249, Y254, E256, F257; Q271’, S274’, E275’) (Fig. 7G). The 22 PomX^NPEP^ residues, including those predicted to interact with PomZ, are highly conserved in PomX orthologs (Fig. 7H); and, the residues in contact distance to PomX^NPEP^ are highly conserved in PomZ orthologs, but not in ParA and MinD orthologs (Fig. 7E; Fig. S11), thus lending further support to the structural model.

In the third step, we modeled dimeric PomZ with two molecules each of PomY^496-682^ and PomX^NPEP^. As in the previous models, the PomZ dimer, α5 and the gD domain in PomY^496-682^, and PomX^NPEP^ were modelled with high confidence based on pLDDT scores (Fig. S12A). Likewise, based on pAE scores, only the two α5s and the two PomX^NPEP^ molecules were modeled with high confidence to interact with the PomZ dimer (Fig. S12A). Both α5s and both PomX^NPEP^ molecules were modeled to bind the PomZ dimer symmetrically in clefts between the two PomZ protomers and within contact-distance to residues from both PomZ protomers; these binding sites are similar, but not identical, to those predicted when PomY^496-682^ and PomX^NPEP^ were modeled separately (Fig. 8A-B). Because the positions of the α5s were slightly shifted, α5 residue V584 was not modeled in contact distance to PomZ (Fig. 8C; cf. Fig. 7C), while M591 and A590 were predicted to be closer to L146, L182 and A147 in PomZ (Fig. 8C; cf. Fig. 7C). Of note, A590 is also highly conserved in PomY orthologs predicted to have an α5 (Fig. 7D) and A147 is highly conserved in PomZ orthologs but not in ParA and MinD orthologs (Fig. 7E; Fig. S11). For PomX^NPEP^, only residues 3-8 were modeled as α-helical (Fig. 8D); however, the PomX^NPEP^ and PomZ residues in contact distance were the same as when PomX^NPEP^ was modeled separately, although the predicted interactions were slightly different (Fig. 8D; cf. Fig. 7G). In agreement with both binding modes of PomX^NPEP^ to the PomZ dimer, a previous Ala-scanning of the 11 charged or hydrophilic residues in PomX^NPEP^ demonstrated that only the two positively charged residues K13 and R15 (Fig. 7H), which are modeled in close proximity to PomZ in both models, are required for PomX activity *in vivo* and AAP activity *in vitro* (27). Conversely, K2, which is also modeled in close proximity to PomZ in both models, when substituted to Ala did not interfere with PomX activity *in vivo*, while AAP activity *in vitro* was not investigated (27).

**Figure 8.**
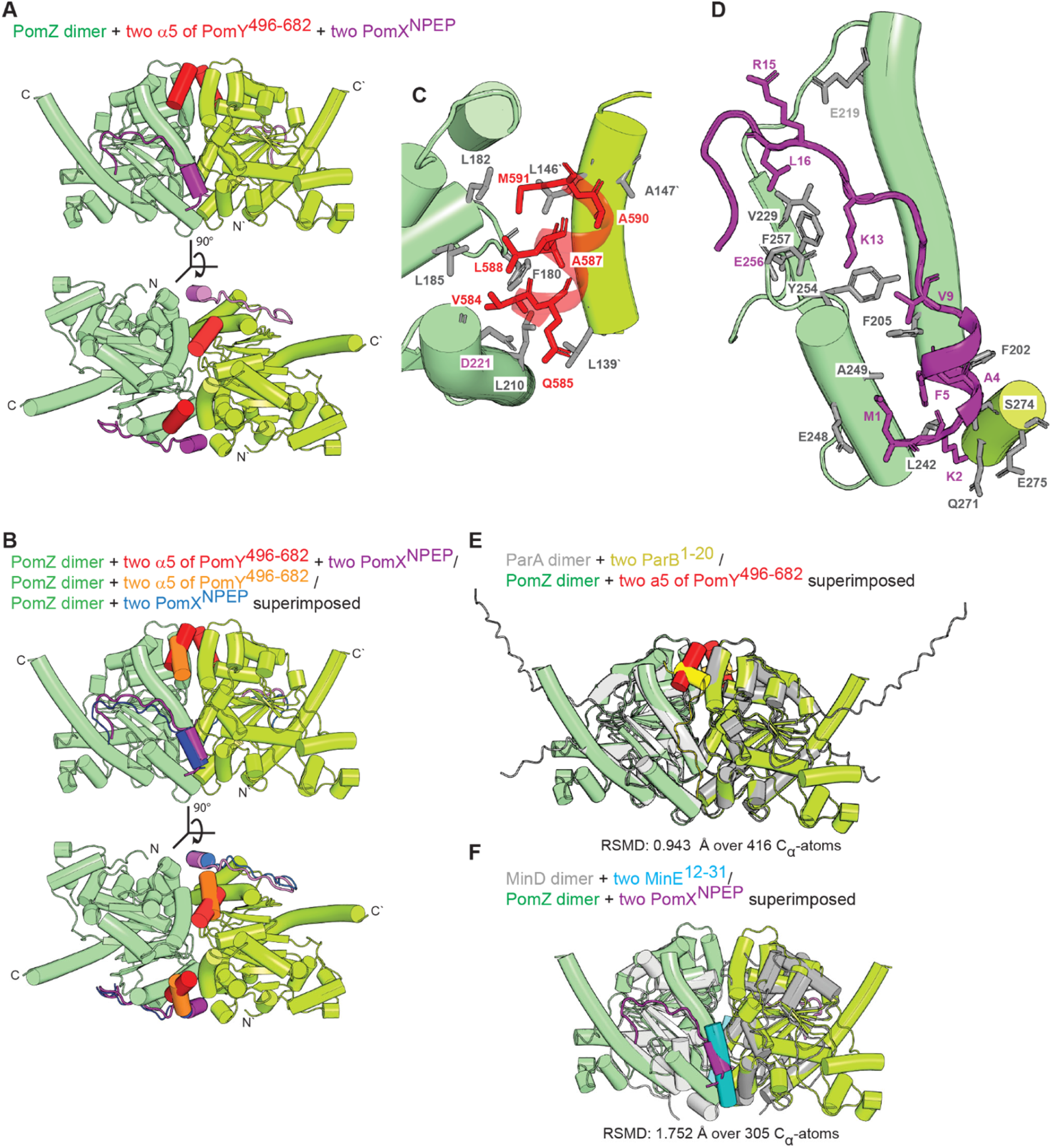
Structural modeling of the PomZ/PomY^496-682^/PomX^NPEP^ interactions and comparison to the ParA/ParB and MinD/MinE interactions. (A) AF3-based structural model of PomZ dimer (green shades) with two copies of α5 of PomY^496-682^ (red) and two molecules of PomX^NPEP^ (purple). The model was generated with PomY^496-682^, but only the high confidence interactions of the two α5s to the PomZ dimer are included. PomZ was modeled with ATP and Mg^2+^ but the ATP molecules and Mg^2+^ are not included for simplicity. Model rank 1 is shown. (B) Comparison of the binding modes of the PomZ dimer (green shades) to α5 of PomY^496-682^ and PomX^NPEP^. The structure of the PomZ dimer with α5 of PomY^496-682^ (red) and PomX^NPEP^ (purple) in combination was structurally aligned with the two structures of the PomZ dimer with either α5 of PomY^496-682^ (orange) or PomX^NPEP^ (blue). (C) Zoom of the predicted interaction of the PomZ dimer and α5 of PomY^496-682^ from the model in A. PomZ residues (grey) and α5 residues (red) predicted to be involved in the interaction are shown in stick representation. (D) Zoom of the predicted interaction of the PomZ dimer and PomX^NPEP^ from the model in A. PomZ residues (grey) and PomX^NPEP^ residues (purple) predicted to be involved in the interaction are shown in stick representation. (E-F) Structural alignment of PomZ dimer (green shades) with two molecules of α5 of PomY^496-682^ (red) and two molecules of PomX^NPEP^ (purple) to ParA dimer (grey shades) with two molecules of ParB^1-20^ (yellow) (E) and MinD dimer (grey shades) with two molecules of MinE^12-31^ (cyan) (F). The ParA/B proteins are those of *M. xanthus* and the MinD/E proteins are those *of E. coli*.

The structural modeling and previous experimental analyses support that PomY^496-682^ *via* α5 and PomX^NPEP^ interact with both protomers at the PomZ dimer interface and would be able to bind separately as well as simultaneously to the PomZ dimer. However, we also note that the PomY^496-682^ interaction *via* α5 may not be the only interaction between PomY and PomZ because PomY^496-623^, which contains α5, is not sufficient to stimulate PomZ ATPase activity (Fig. 4F).

### Comparison of the binding sites of α5 of PomY^496-682^ and PomX^NPEP^ in the PomZ dimer to those of ParB in the ParA dimer and MinE in the MinD dimer

The most detailed analysis of the ParA/ParB interaction based on the *M. xanthus* proteins revealed that two copies of a partially α-helical N-terminal 20-residues peptide of ParB (ParB^1-20^), which is required and sufficient for AAP function, bind symmetrically to the ParA dimer in a cleft at the dimer interface making contacts to both protomers (26) (Fig. S12B) (26). Binding is driven by hydrophobic interactions between ParB^1-20^ and the ParA dimer, while R13 in ParB^1-20^ is essential for stimulation of ParA ATPase activity (26). In the *E. coli* MinD/MinE system, two copies of an α-helical 20-residue N-terminal peptide of MinE (MinE^12-31^) bind symmetrically to the MinD dimer interface contacting both protomers in the solved structure (Fig. S12C) (24, 35). Binding is driven by a mixture of hydrophobic and hydrophilic interactions, and a positively charged residue (R21) in MinE^12-31^ is essential for AAP activity (24, 35, 42). The ParB^1-20^ and MinE^12-31^ binding sites are at different positions at the ParA/MinD dimer interface (Fig. S12B-C). Interestingly, the predicted binding site of α5 of PomY in the PomZ dimer partially overlaps with the ParB^1-20^ binding site in the ParA dimer (Fig. 8E). Most of the predicted interactions between PomZ/α5 and ParA/ParB^1-20^ are hydrophobic (Fig. 7C, Fig. 8C) (This study; (26)). However, α5 of PomY does not contain the positively charged residue (R13), which is essential in ParB/ParB^1-20^ for ATPase stimulation (26). Remarkably, the binding sites of PomX^NPEP^ and MinE^12-31^ in the PomZ and MinD dimer, respectively also partially overlap (Fig. 8F). In both cases, binding is driven by a mixture of hydrophobic and hydrophilic interactions (Fig. 8C) and both peptides contain positively charged residue(s) important for AAP function (K13 R15 in PomX^NPEP^ and R21 in MinE^12-31^).

These analyses suggest that PomY and PomX use a combination of the two distinct and experimentally verified binding sites of ParB and MinE to their partner ATPases to synergistically activate PomZ ATPase activity.

## Discussion

The PomX/Y/Z system in *M. xanthus* is important for positioning the division site at midcell. As opposed to other systems incorporating a ParA/MinD ATPase, the PomX/Y/Z system incorporates two AAPs, PomX and PomY. While prior work established that the AAP activity of PomX resides in a positively charged 22-residues N-terminal peptide, the mechanism of PomY in the activation of PomZ activity remained unclear (27). Here, using *in vitro* analyses, we map the PomY AAP activity to the C-terminal PomY^496-682^ region and show that PomY, similarly to PomX (27), interacts with the ATP-bound PomZ dimer. These observations strongly suggest that PomX and PomY both directly stimulate PomZ ATPase activity rather than nucleotide exchange. We propose that by restricting the interaction of PomX and PomY to the ATP-bound PomZ dimer that binds non-specifically to DNA, the PomX/Y assembly is efficiently restricted to interact with the nucleoid-bound PomZ dimer *in vivo*.

*In vivo*, the PomY^Δ496-682^ variant that lacks AAP activity formed clusters as efficiently as WT PomY, documenting that the interaction between PomY^Δ496-682^ and PomX was not compromised. However, these clusters were less frequently at midcell than those of WT PomY. In agreement with PomY^496-682^ having AAP activity and interacting with PomZ, PomZ was less efficiently recruited to the PomX/Y assembly in the PomY AAP^-^ mutant. Altogether, these observations strongly suggest that the PomY-dependent stimulation of PomZ ATPase activity is important for the translocation of the PomX/Y/Z assembly to midcell. Importantly, these observations parallel those of PomX AAP^-^ mutants, which are also compromised in localizing the PomX/Y/Z assembly to midcell and in recruiting PomZ to the PomX/Y assembly (27). Collectively, these observations suggest that stimulation of PomZ ATPase activity by both PomX and PomY is essential for efficient translocation of the PomX/Y/Z assembly to midcell.

In agreement with the localization of the PomY^Δ496-682^ AAP^-^ variant, cell division constrictions occurred less frequently at midcell than in WT. Interestingly, constrictions were also formed at a reduced frequency, and occasionally they did not occur over the PomY^Δ496-682^ cluster. Thus, the PomY AAP^-^ mutant is compromised in stimulating cell division. By comparison, (i) in a Δ*pomY* mutant, constrictions are formed at a reduced frequency, away from midcell, and essentially none of these constrictions occurs over the Pom cluster; (ii) in a Δ*pomZ* mutant, constrictions are formed at a reduced frequency and away from midcell, but constrictions occur over the Pom cluster; and (iii) in PomX AAP mutants, constrictions are formed at WT frequencies and away from midcell, but constrictions occur over the Pom cluster. These comparisons indicate that mislocalization of the Pom cluster away from midcell is not responsible for the constriction defect observed in the PomY AAP mutant. Rather these comparisons support a model in which the formation of a cell division constriction occurs in a two-stage process. In the first stage, FtsZ is recruited to the nascent division site, PomY is key to this stage, while PomZ and PomX AAP activity are dispensable. In the second stage, FtsZ polymerization is stimulated; PomY as well as PomZ are key to completion of this stage, while PomX AAP activity is dispensable. Based on the phenotype of the PomY^Δ496-682^ AAP^-^ mutant, we suggest that it retains partial PomY activity and is able to recruit FtsZ to the nascent division site almost as efficiently as WT PomY, but it is deficient in supporting FtsZ polymerization to form the Z-ring and, thus, initiate cell division. In agreement with this model, PomY biomolecular condensates *in vitro* enrich FtsZ and stimulate the GTP-dependent polymerization of FtsZ (6); and, based on Y2H analyses, PomZ also interacts with FtsZ (7). Therefore, in the future it will not only be interesting to test *in vitro* which PomY part(s) interacts with FtsZ and stimulates the GTP-dependent polymerization but also to test whether PomZ and its ATPase activity are important for the GTP-dependent FtsZ polymerization.

Structural modeling of the PomY^496-682^ region suggests that it is largely an IDR but also contains the nine-residue α5 and the gD domain. The gD domain adopts a wHTH fold, but sequence analyses and experiments suggest that it neither binds Z-DNA nor DNA. Importantly, neither the IDR with α5 nor the gD domain is sufficient for PomY AAP activity, documenting that determinants in both subregions are required for PomY AAP activity. Surprisingly, AF3-based structural modeling only supported an interaction between α5 and the PomZ dimer with high confidence. By contrast, an interaction between the gD domain and the PomZ dimer or other parts of the PomY^496-682^ region was not supported by the structural modeling. The structural modeling suggested that two α5 copies bind the PomZ dimer symmetrically in a mostly hydrophobic cleft between the two protomers and making contact to residues in both protomers, thus explaining why PomY preferentially interacts with the ATP-bound PomZ dimer. Interestingly, the α5 binding site partially overlaps with the hydrophobic binding site of the ParB^1-20^ peptide, which is required and sufficient for ParB AAP activity, in the ParA dimer (26). However, ParB^1-20^ also interacts with the ParA dimer through a positively charged residue (R13), which is essential for AAP activity but not for binding (26). This residue has been proposed to induce a reconfiguration of active-site residues in ParA that stimulates ATP hydrolysis (26). Notably, α5 does not contain a positively charged residue, possibly explaining why the 496-623 region of PomY, which contains α5, is not sufficient for AAP activity. While our results suggests that the α5/PomZ interaction is critical for PomY AAP activity, it is important to acknowledge that this is likely only one of the interactions between the PomY^496-682^ region and the PomZ dimer required for AAP activity. Altogether, based on experimental data and structural modeling, we propose that PomY uses a non-canonical AAP mechanism distinct from those of other well-characterized AAPs of ParA/MinD ATPases, in which a positively charged N-terminal peptide is critical for AAP activity. We propose that α5 mediates binding to the PomZ dimer, whereas other parts of the PomY^496-682^ region contribute to AAP function through an unknown mechanism. Having mapped AAP activity to the PomY^496-682^ region and identified a potential role for α5 in this process, it will be important in future work to address how the remaining parts of this region contribute to AAP activity.

Structural modeling supported with high confidence that two PomX^NPEP^ molecules also bind symmetrically to the PomZ dimer in a cleft between the two protomers and making contact to residues in both protomers, again explaining why PomX preferentially interacts with the ATP-bound PomZ dimer. Interestingly, the PomX^NPEP^ binding site on the PomZ dimer partially overlaps with that of the MinE^12-31^ peptide on the MinD dimer (35). PomX^NPEP^ AAP activity depends on the two positively charged residues K13 and R15, which face towards PomZ in the structural model, and in the MinE^12-31^ peptide, the positively charged residue R21 is essential for AAP activity (24). In the case of the MinE^12-31^ peptide, it has been suggested that its binding to the MinD dimer causes a reconfiguration of residues in the MinD active site and, thus, increased ATPase activity (24). We suggest that PomX^NPEP^ functions analogously to the MinE^12-31^ peptide, and that its binding to the PomZ dimer induces a reconfiguration of the PomZ active site, resulting in increased PomZ ATPase activity.

PomY^496-682^ and non-specific DNA as well as PomX^NPEP^ and non-specific DNA (27) synergistically activate PomZ ATPase activity. In addition, PomY^496-682^ and PomX^NPEP^together synergistically stimulate PomZ ATPase activity. The α5 of PomY^496-682^ and PomX^NPEP^ bind to non-overlapping sites on the PomZ dimer, both located away from its DNA-binding patch. This spatial arrangement provides an explanation for how DNA and the two peptides can simultaneously bind the PomZ dimer and synergistically stimulate ATPase activity. Recently, it was shown that the ParB^1-20^ peptide and non-specific DNA synergistically activate ParA ATPase activity by causing a reconfiguration of active site residues (26). By the same token, we propose that PomY^496-682^, PomX^NPEP^ and non-specific DNA synergistically activate PomZ ATPase activity by binding to distinct sites on the dimer, thereby inducing reconfigurations of its active-site residues.

## Supporting information

All Supplemenary Information

## Acknowledgment

We thank Dominik Schumacher for plasmids and helpful discussions. This work was generously supported by the Max Planck Society (to LS-A).

## Availability of data and materials

The data supporting the findings of this study are all included in the manuscript and its supplementary file. All materials are available from the corresponding author.

## Materials and Methods

### *M. xanthus* and *E. coli* strains and growth

All *M. xanthus* strains used in this study are derivatives of the WT strain DK1622 (43) and are listed in Table S1. Plasmids and oligonucleotides used are listed in Table S2 and S3, respectively. Plasmid construction is described in Table S4. All *M. xanthus* strains used contain an in-frame deletion of *mglA* and are non-motile (44, 45). In-frame deletions in *M. xanthus* were constructed *via* two-step homologous recombination as described (46). Plasmids for complementation experiments were integrated in a single copy by site-specific recombination at the Mx8 *attB* site. Point mutations were introduced using the QuikChange II XL Site-Directed Mutagenesis Kit (Agilent). All plasmids were verified by DNA sequencing, and all strains were verified by PCR. *M. xanthus* cultures were grown at 32°C in 1% CTT broth (1% [w/vol] Bacto casitone, 10 mM Tris-HCl [pH 8.0], 1 mM K_2_HPO_4_/KH_2_PO_4_ [pH 7.6], 8 mM MgSO_4_) or on 1.5% agar supplemented with 1% CTT and kanamycin (50 μg mL^−1^), oxytetracycline (10 μg mL^−1^) or gentamycin (10 μg mL^−1^) when appropriate (47). Growth was measured as an increase in optical density (OD) at 550 nm. Plasmids were propagated in *E. coli* NEB Turbo (New England Biolabs) (F’ *proA^+^B^+^ lacI^q^ ΔlacZM15/fhuA2 Δ(lac-proAB) glnV galK16 galE15 R(zgb-210::Tn10)* Tet^S^ *endA1 thi-1 Δ(hsdS-mcrB)5*) in lysogeny broth (LB) (48) supplemented with ampicillin (100 µg mL^-1^), kanamycin (50 μg mL^−1^) or tetracycline (20 μg mL^−1^) when required.

### Gal4-based Yeast Two hybrid (Y2H) Assay

Yeast two hybrid assays were performed as described by the manufacturer (Clontech). Briefly, genes of interest were fused to the Gal4AD fragment (activation domain) or the Gal4BD fragment (DNA-binding domain). Plasmids were co-transformed into yeast strain AH109. Transformants were selected on SD/-Leu/-Trp 2% agar (Takara) for the presence of both plasmids. Three independent clones were resuspended in SD/-Leu/-Trp medium and grown in suspension culture for three doubling times at 30°C. OD_600_ was adjusted to 0.5 and 4μl aliquots of cells were placed on SD/-Leu/-Trp/-His (low stringency), or SD/-Leu/-Trp/-His + 10mM 3-amino-1,2,4-triazole (3AT) (high stringency) selective agar. Plates were incubated at 30°C for 120 hrs. Additionally each plasmid containing a gene of interest was co-transformed with an empty vector only expressing the Gal4AD or Gal4BD fragment. In an experiment, all strains were spotted with all the controls on the same selective agar plate.

### Fluorescence microscopy

Fluorescence microscopy was performed as described (7). Briefly, exponentially growing cultures were transferred to slides with a thin 1.0% agarose pad (SeaKem LE agarose, Cambrex) with TPM buffer (10 mM Tris-HCl [pH 7.6], 1 mM KH_2_PO_4_ [pH 7.6], 8 mM MgSO_4_), covered with a coverslip and imaged using a temperature-controlled Leica DMi8 inverted microscope with a 100× HCX PL FLUOTAR objective at 32°C. Phase-contrast and fluorescence images were recorded with a Leica K8 camera using the LASX software (Leica Microsystems). Image processing was performed with Metamorph v 7.5 (Molecular Devices). For image analysis, cellular outlines were obtained from phase-contrast images using Oufti (49) and manually corrected if necessary. Fluorescence microscopy image analysis was performed with a custom-made Matlab script (Matlab R2018a, MathWorks) as described (27). Constrictions were identified manually.

### Immunoblot analysis

For immunoblot analyses, samples were prepared by harvesting *M. xanthus* cells from exponentially growing suspension cultures, followed by resuspension of the cell pellet in SDS lysis buffer (60 mM Tris-HCl [pH 6.8], 2% SDS [w/vol], 10% glycerol [vol/vol], 0.1 M dithiothreitol, 0.005% bromophenol blue [w/vol]). Proteins of an equal amount of cells were loaded per sample. Immunoblot analyses were performed as described (50) with rabbit polyclonal α-PomY (1:15000) (7), α-PomZ (1:10000) (8), α-PilC (1:3000) (51), or α-mCh (1:10000; Biovision) primary antibodies together with horseradish-conjugated goat α-rabbit immunoglobulin G (Sigma-Aldrich) as secondary antibody (1:25000). Blots were developed using Luminata Forte Western HRP Substrate (Millipore) and visualized using an Amersham ImageQuant 800 (Cytiva).

### Protein purification

His_6_-PomZ was purified from *E. coli* containing the plasmid pKA3 as described (8). Briefly, Rosetta 2 [F-*ompT hsdSB* (rB-mB-) *gal dcm*] (Novagen) containing pKA3 and pRARE2 was grown in 2×YT medium supplemented with 0.2% glucose and 100 µg ml^-1^ ampicillin to mid-exponential phase at 37°C. Overexpression was induced with 0.5 mM IPTG for two hr. Cells were harvested at 4°C for 10 min at 4,400× *g*, washed and resuspended in buffer DB (50 mM HEPES/NaOH [pH 7.2], 50 mM KCl, 0.1mM EDTA, 10% glycerol, 1 mM β-mercaptoethanol) supplemented with 5 mM MgCl_2_. After two passages through a French press (16,000 psi), the cell suspension was centrifuged at 4°C for 30 min at 30,000× *g*. Soluble His_6_-PomZ was purified on a 5 ml HiTrap Chelating HP column (GE Healthcare). Fractions containing His_6_-PomZ were pooled and dialyzed against buffer DB supplemented with 5 mM MgCl_2_. After centrifugation at 4°C for 30 min at 30,000× *g*, the supernatant was applied to a Hiload 16/60 Superdex 200pg size exclusion column (1.6 × 60 cm, GE Healthcare) equilibrated with buffer DB. Fractions containing His_6_-PomZ were pooled, frozen in liquid N_2_ and stored at -80°C.

PomY-mCh-His_6_ (pAH194) and all variants (PomY-His_6_ (pDS3), PomY^CC^-mCh-His_6_ (pAH196), PomY^HEAT^-mCh-His_6_ (pAH197), PomY^mIDR^-mCh-His_6_ (pAH198), PomY^CC_HEAT^-mCh-His_6_ (pAH187), PomY^HEAT_mIDR^-mCh-His_6_ (pAH195), PomY^CC-mIDR^-mCh-His_6_ (pAH217), PomY^Δ438-488^-mCh-His_6_ (pPK57), PomY^Δ496-623^-mCh-His_6_ (pPK67), PomY^Δ624-682^-mCh-His_6_ (pPK68), His_6_-Sumo-PomY^mIDR^-mCh (pPK83), His_6_-Sumo-PomY^496-623^-mCh (pPK84), His_6_-Sumo-PomY^624-682^-mCh (pPK85), His_6_-Sumo-PomY^496-682^-mCh (pPK86)) were purified as described (6). Briefly, to purify PomY-mCh-His_6_ and variants, the corresponding plasmids were propagated in *E. coli* ArcticExpress(DE3) RP cells (*E. coli B F– ompT hsdS*(*rB– mB-*) *dcm+ Tetr gal λ(*DE3) *endA Hte* [*cpn10 cpn60 Gentr*] [*argU proL Strr*]) (Agilent Technologies), grown in 2×YT medium with 50 µg ml^-1^ kanamycin at 30°C to an OD_600_ of 0.6–0.7. Protein expression was induced with 1 mM IPTG for 16 h at 18°C. Cells were harvested by centrifugation at 5,000× *g* for 20 min at 4 °C and washed in lysis buffer 1 (50 mM NaH_2_PO_4_, 300 mM NaCl, 10 mM imidazole, 20 mM β-mercaptoethanol, [pH 8.0]). Cells were lysed in 50ml lysis buffer 2 (lysis buffer 1, 100 µg ml^-1^ phenylmethylsulfonyl fluoride (PMSF), 1× complete protease inhibitor (Roche Diagnostics GmbH), 10 U ml^-1^ benzonase, 0.1% Triton X-100, [pH 8.0]) by three rounds of sonication for 5 min with a UP200St ultrasonic processor (pulse 80%, amplitude 80%) (Hielscher) on ice. Cell debris was removed by centrifugation (4,150× *g* for 45 min at 4 °C) and filtration with a 0.45-μm sterile filter (Millipore Merck). PomY-mCh-His_6_ and variants were purified with a 5 ml HiTrap Chelating HP (Cytiva, 17040901) column preloaded with NiSO_4_ and equilibrated with lysis buffer 1. The column was washed with 20 CV (column volumes) wash buffer 1 (lysis buffer 1, 20 mM imidazole) and five CV wash buffer 2 (lysis buffer 1, 50 mM imidazole). Protein was eluted with elution buffer (lysis buffer 1, 300 mM imidazole). Protein was then applied to a HiPrep 26/60 Sephacryl S-400 HR size exclusion column (Cytiva; 28935605) equilibrated in dialysis buffer (50 mM HEPES/NaOH, 1M KCl, 0.1 mM EDTA, 20 mM β-mercaptoethanol, 10% (v/v) glycerol, [pH 7.2]). Fractions were dialysed in dialysis buffer supplemented with 50mM KCl and snap-frozen in liquid nitrogen and stored at −80 °C until use. Protein concentration was determined immediately before use using Protein Assay Dye Reagent Concentrate (BioRad).

### ATPase assay

ATP hydrolysis was determined using a 96-well NADH-coupled enzymatic assay (52) with modifications. Protein concentration was determined using Protein Assay Dye Reagent Concentrate (BioRad). Assays were performed in reaction buffer (50 mM HEPES/NaOH [pH 7.2], 50 mM KCl, 10 mM MgCl_2_) with 0.5 mM nicotinamide adenine dinucleotide (NADH) and 2 mM phosphoenolpyruvate and 3 µl of a pyruvate kinase/lactate dehydrogenase mix (PYK/LDH; Sigma). PomX^NPEP^ peptide (MKKAFEQNVSRAKPRLRLGALT) was purchased from Thermo Scientific as previously described (27). Herring sperm DNA was added to a final concentration of 60 µg ml^-1^ unless otherwise stated. Buffer was pre-mixed with proteins in low-binding microtubes (Sarstedt) on ice. To correct for glycerol in the assays, dialysis buffer was added if necessary. A total of 100 µl mixtures were transferred into transparent UV-STAR µCLEAR 96-well microplates (Greiner bio-one). The reaction was started by the addition of 1 mM ATP. Measurements were performed in an infinite M200PRO (Tecan) for 2 hr in 30 s intervals at 32°C at 340 nm wavelength. To account for background by spontaneous ATP hydrolysis and UV-induced NADH decomposition, all assays were performed without the addition of His_6_-PomZ and measurements were subtracted. The light path was determined experimentally with known NADH concentrations to be 0.248 cm. The extinction coefficient of NADH ε340 = 6220 M^−1^cm^−1^ was used.

### Biolayer interferometry (BLI)

BLI analyses were performed with a BLItz® system using High Precision Streptavidin 2.0 (SAX2) Biosensors (Sartorius). A 236 bp DNA fragment was PCR-amplified from *M. xanthus* chromosomal DNA with two primers (cluster3 prom 1 bioteg and cluster3 prom 2 bioteg) each carrying a 5’-biotin-triethylene glycol (TEG) group. After purification with a Monarch Spin PCR and DNA Cleanup kit (New England Biolabs), the double-biotinylated fragment (100 nM) was immobilized on the biosensor in BLI binding buffer (50 mM HEPES/NaOH [pH 7.2], 50 mM KCl, 0.1mM EDTA, 10% [v/v] glycerol, 10 mM MgCl_2_, 1 mM β-mercaptoethanol, 0.05% [v/v] Tween 20), and a stable baseline established. The relevant proteins were mixed in BLI binding buffer supplemented with 1mM ATP on ice for 10 min, and loaded on the sensor. At the end of the association step, the biosensor was transferred into protein-free BLI binding buffer with ATP to follow the dissociation kinetics. The K_D_ was calculated in GraphPad Prism with a One-site specific binding model.

### Bioinformatics

Gene and protein sequences were obtained from the databases of KEGG (53) or NCBI (54). PomY orthologs were identified in a best-best hit reciprocal BlastP analysis from fully-sequenced genomes of Myxobacteria (55). The similarity and identity of proteins were calculated from pairwise sequence alignments with EMBOSS Needle (56). Multiple sequence alignments were created with MAFFT (56) and further edited with Bioedit (https://bioedit.software.informer.com/7.2/). Consensus sequences of multiple sequence protein alignments were created with Weblogo 3 (57). Protein domains were predicted using SMART (58), Interpro (59), MyHits (60), and the structural AF3 predictions. Protein hydropathy was analyzed with Protscale (61) with the parameters of Kyte & Doolittle (62) and a window size of three. Protein charge was analyzed with EMBOSS:charge (https://www.bioinformatics.nl/cgi-bin/emboss/charge).

All structural predictions were performed with AlphaFold3 (AF3) via the AlphaFold server (https://alphafoldserver.com/) with default settings (37). To evaluate AF3-generated models, predicted local distance difference test (pLDDT) and predicted alignment error (pAE) graphs of five models were made using a custom-made Matlab R2020a (The MathWorks) script. These models were ranked based on combined pLDDT, pAE, and predicted (interface) template modeling score ((i)pTM) scores, with the best-ranked models used for further analysis and presentation. Per-residue model confidence was estimated based on pLDDT scores (>90, high accuracy; 70–90, generally good accuracy; 50–70, low accuracy; <50, should not be interpreted) (63). The relative positioning of residues was validated by assessing pAE, measured in Å. The lower the pAE score, the higher the accuracy of the relative position of residue pairs and, consequently, the relative position of domains/subunits/proteins (63). pTM scores were used to assess how well AF3 predicted the overall structure of a complex. Generally, a pTM score >0.50 indicates that a predicted structure is similar to the true structure and lower pTM scores indicate a likely incorrect structure (64). ipTM scores (64) were used to evaluate interface accuracy in multimeric models. Generally, an ipTM score >0.80 indicates an accurate interface, scores between 0.60 and 0.80 represent a gray zone where predictions may or may not be correct, and scores below 0.60 indicate a failed prediction (37, 65). PyMOL (The PyMOL Molecular Graphics System, Version 2.4.1 Schrödinger, LLC) was used to analyze and visualize the structural models. For generating models colored based on pLDDT values, a custom command line was used (spectrum b, red_yellow_green_cyan_blue, minimum = 50, maximum = 90). Structural superimpositions were performed using the “super” method within the PyMOL that was also used for calculating Root Mean Square Deviation (RMSD). The electrostatic surface potential was visualized using the Protein Contact Potential tool implemented in PyMOL. Alignment plugin with default settings. Foldseek (38) was used to identify structural homologs in the PDB.

The coordinates of all structural models generated in this study have been deposited in the Edmond research data repository (https://doi.org/10.17617/3.HEWOMR).

### Statistics

The mean and standard deviation (STDEV) were calculated with Excel 2016 or GraphPad Prism. Localization patterns from fluorescence microscopy data were quantified based on the indicated n-values per strain. Curve fitting was performed with GraphPad Prism. All experiments were performed at least in three biological replicates.

