## Supplementary material for "Molecular insights into the dual activation of PomZ, a ParA/MinD P-loop ATPase, by the two ATPase activating proteins PomX and PomY": All Supplemenary Information

**This file contains:**

- Supplementary Figures 1-12
- Supplementary Tables 1-4
- Supplementary References

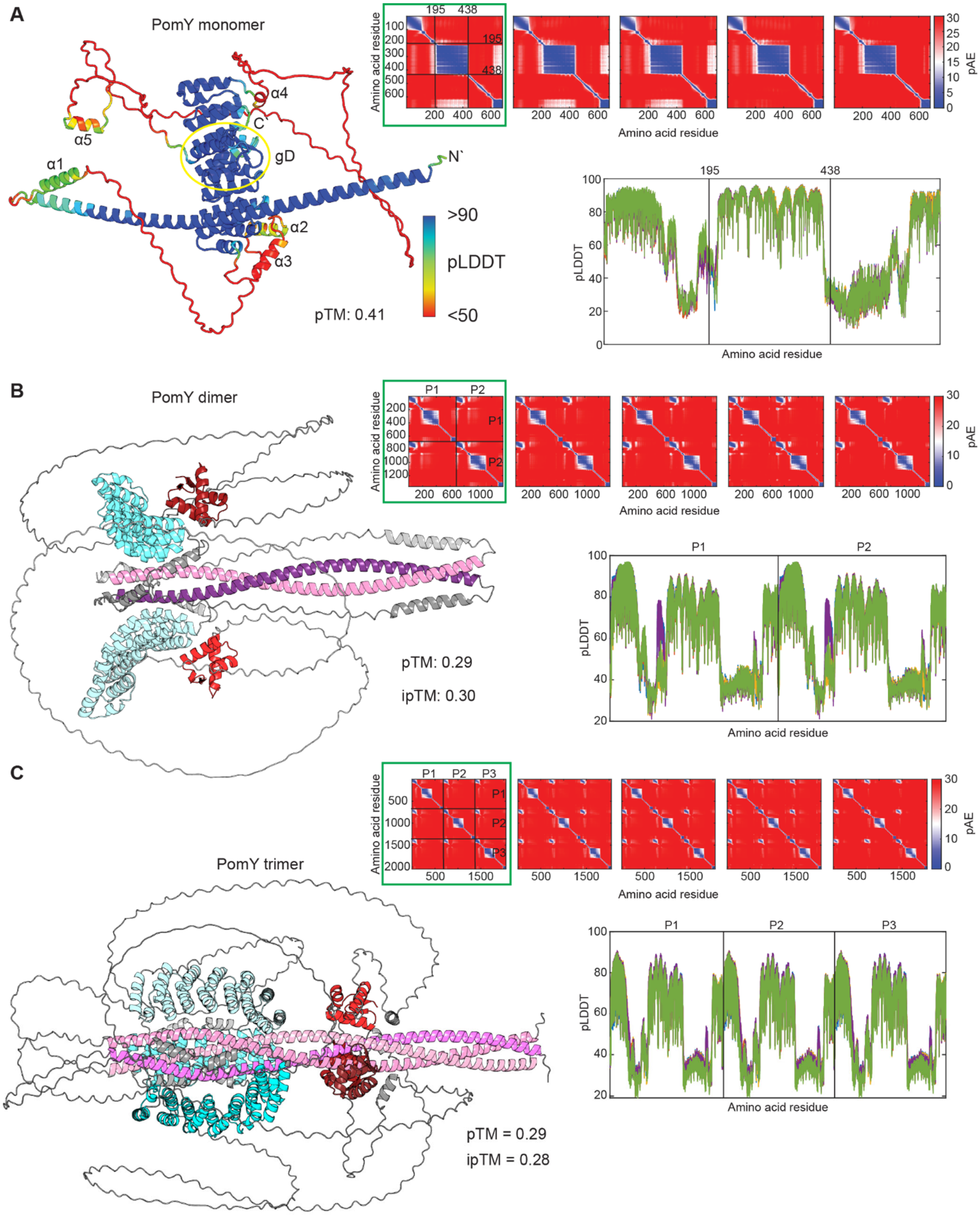


<u>Figure S1. AlphaFold3-based structural models of PomY.</u>

(A) Left panel, model rank 1 of PomY monomer colored according to pLDDT score. The gD domain is indicated by a yellow circle. Right panels, pAE and pLDDT plots of the five models

generated. Black lines indicate truncation points described in Fig. 1A and Fig. 2B. Model rank 1 (highlighted by a green box) was selected for further analyses.
(B-C) Left panel, model rank 1 of PomY dimer (B) and trimer (C). Domains are colored as in Fig. 1A. Right panels in, pAE and pLDDT plots of the five models generated. Black lines separate the two (B) and three (C) protomers. Model rank 1 (highlighted by a green box) was selected for further analysis. Note that the pTM and ipTM scores for both the dimer and trimer are below 0.50 (Fig. S1B-C). Because the pTM and ipTM scores reflect the confidence of the entire structure, the overall confidence of these structures is low due to the large proportion IDRs.

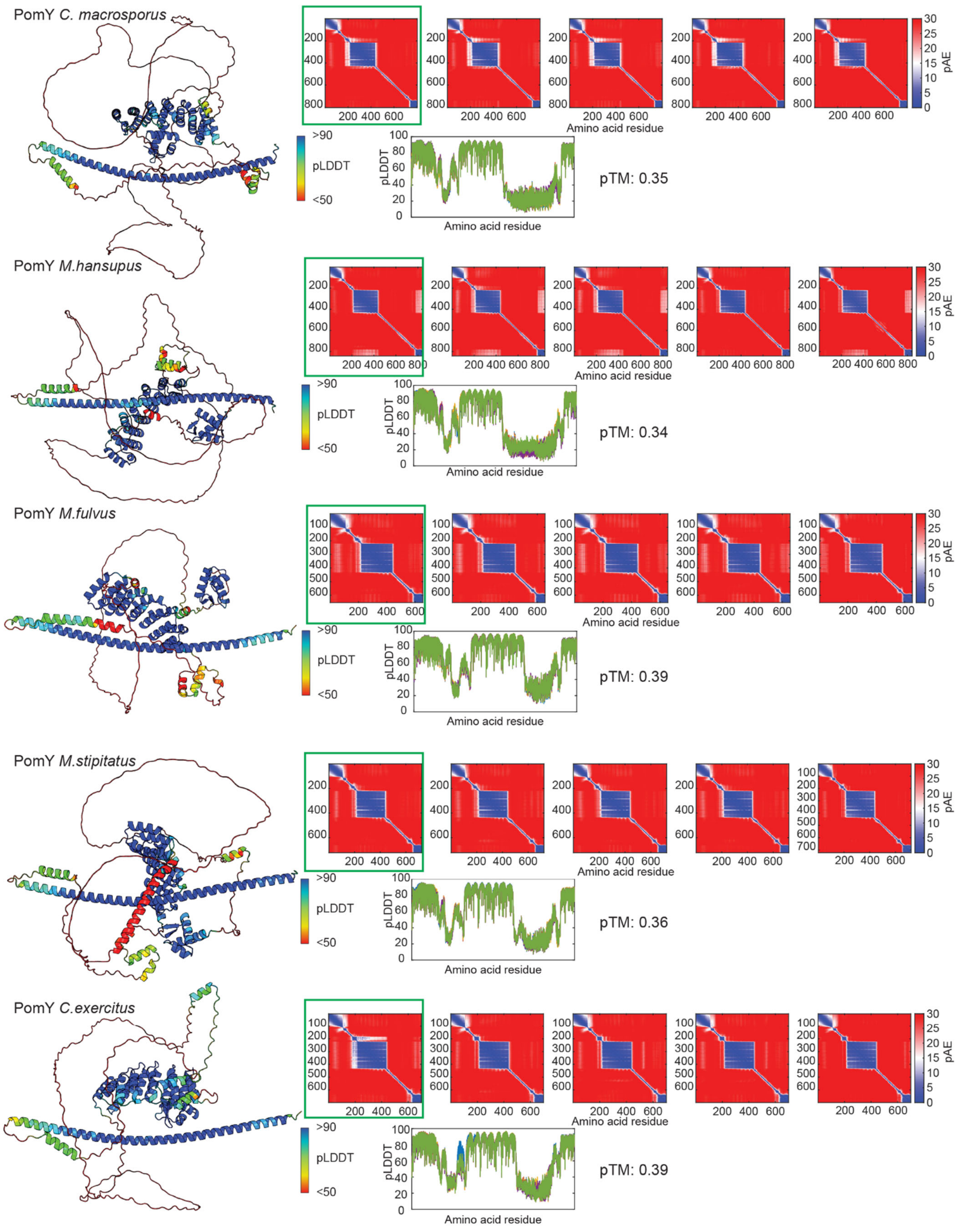


Figure S2. AlphaFold3-based structural models of monomeric PomY orthologs.

Left panels, model rank 1 of the indicated PomY ortholog colored according to pLDDT scores. Right panels, pAE and pLDDT plots of the five models generated. Model rank 1 (highlighted by a green box) was selected for further analysis. Note that the models are of low confidence based on the pTM scores. The large proportion of IDRs, which have low pLDDT scores, causes these low scores.

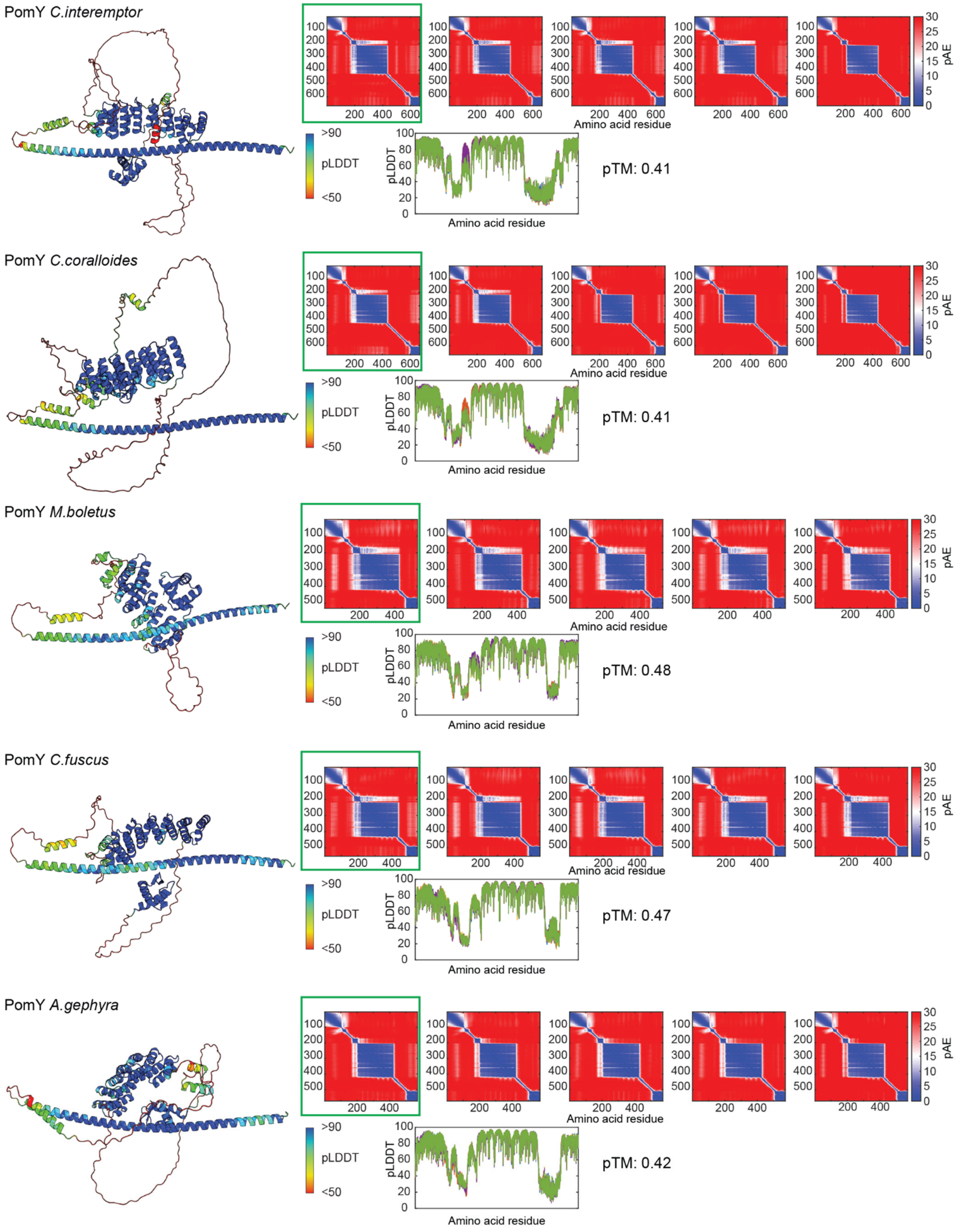


Figure S3. AlphaFold3-based structural models of monomeric PomY orthologs.
Left panels, model rank 1 of the indicated PomY ortholog colored according to pLDDT scores. Right panels, pAE and pLDDT plots of the five models generated. Model rank 1

(highlighted by a green box) was selected for further analysis. Note that the models are of low confidence based on the pTM scores. The large proportion of IDRs, which have low pLDDT scores, causes these low scores.

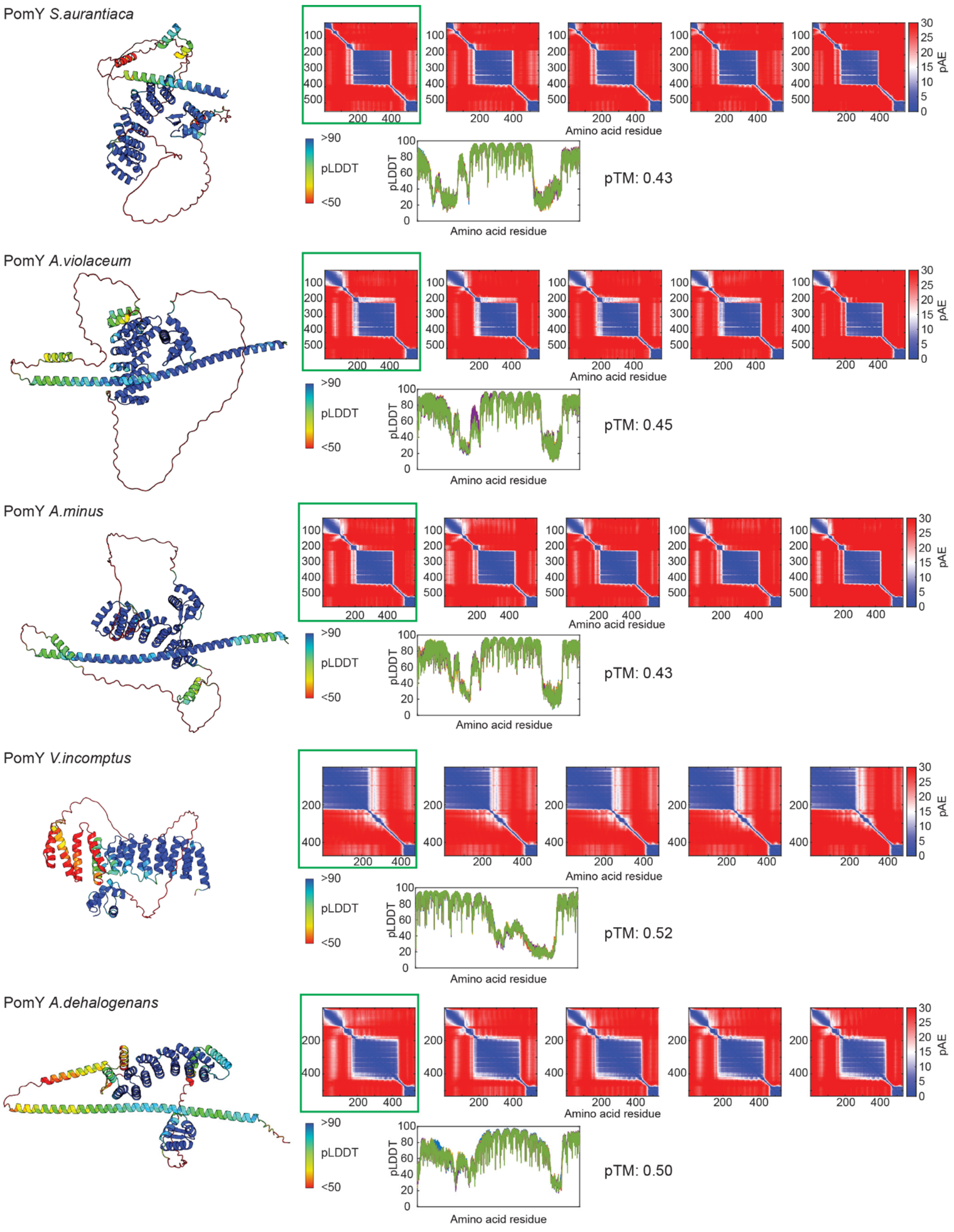


<u>Figure S4. AlphaFold3-based structural models of monomeric PomY orthologs.</u>
Left panels, model rank 1 of the indicated PomY ortholog colored according to pLDDT scores. Right panels, pAE and pLDDT plots of the five models generated. Model rank 1

(highlighted by a green box) was selected for further analysis. Note that the models are of low confidence based on the pTM scores. The large proportion of IDRs, which have low pLDDT scores, causes these low scores.

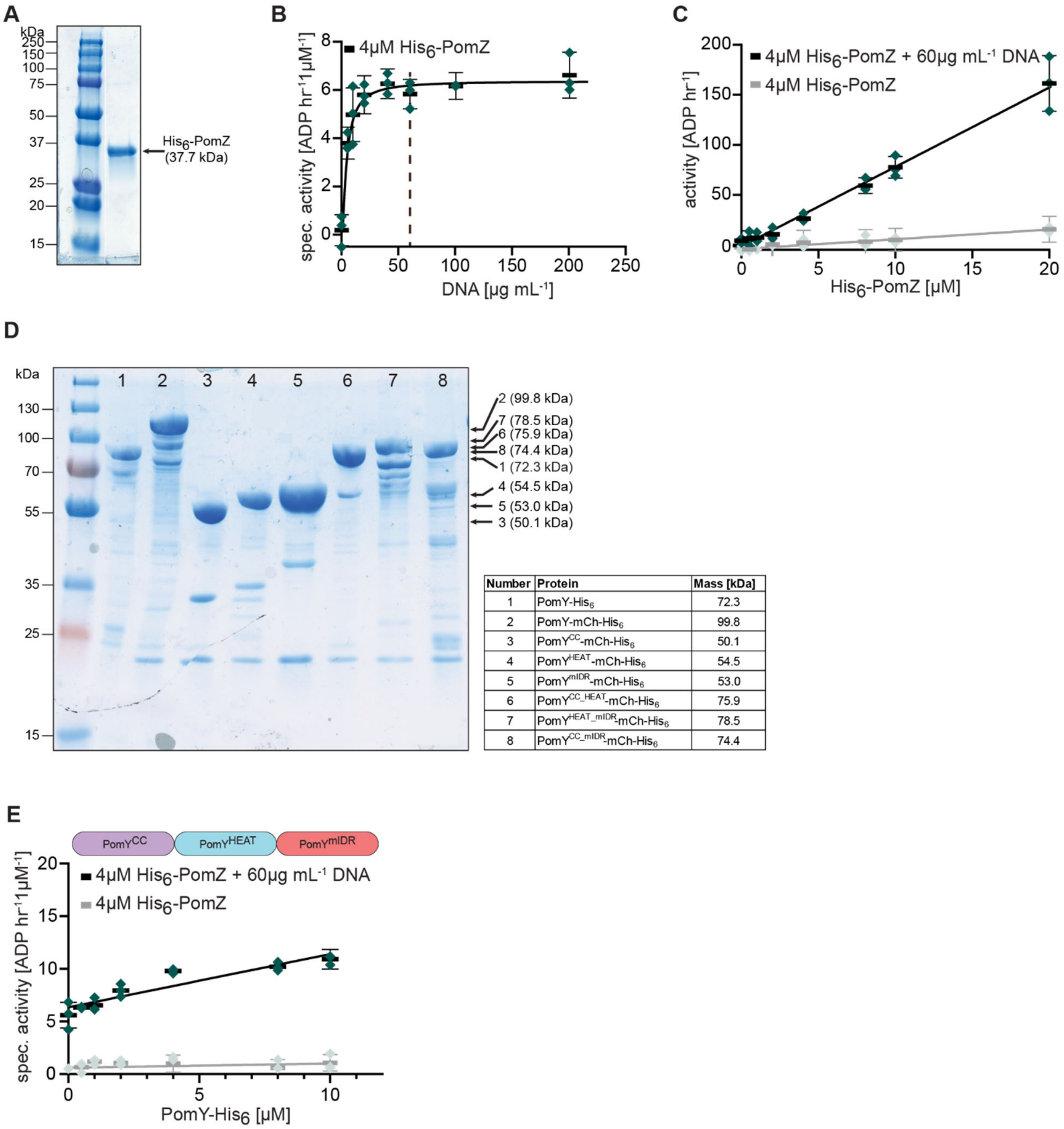


| Number | Protein | Mass [kDa] |
|---|---|---|
| 1 | PomY-His$_6$ | 72.3 |
| 2 | PomY-mCh-His$_6$ | 99.8 |
| 3 | PomY$^{CC}$-mCh-His$_6$ | 50.1 |
| 4 | PomY$^{HEAT}$-mCh-His$_6$ | 54.5 |
| 5 | PomY$^{mIDR}$-mCh-His$_6$ | 53.0 |
| 6 | PomY$^{CC_HEAT}$-mCh-His$_6$ | 75.9 |
| 7 | PomY$^{HEAT_mIDR}$-mCh-His$_6$ | 78.5 |
| 8 | PomY$^{CC_mIDR}$-mCh-His$_6$ | 74.4 |

<u>Figure S5. Purification and analysis of PomZ and PomY variants.</u>
(A) SDS-PAGE analysis of purified His$_6$-PomZ. Molecular size markers are shown on the left and purified His$_6$-PomZ including calculated MW on the right.
(B) His$_6$-PomZ ATPase activity. Double stranded herring sperm DNA (DNA) was added as indicated. The assay was performed as described in Fig. 2A. Green points represent one biological replicate, which was calculated as the mean of three technical replicates. Black bars and error bars, mean of three biological replicates and STDEV. Spontaneous ATP hydrolysis and NADH consumption were accounted for by subtracting the measurements in the absence of His$_6$-PomZ.
(C) His$_6$-PomZ ATPase activity. His$_6$-PomZ and DNA were added as indicated. The assay was performed and analysed as in (B).
(D) SDS-PAGE analysis of purified PomY variants. Left lane, molecular size markers with indicated masses. Each protein was assigned a number, which is detailed in the table on the right. As previously described, PomY-His$_6$ as well as PomY-mCh migrated aberrantly in

SDS-PAGE (1, 2). Proteins were purified as mCh-His$_6$-tagged proteins; in the main text, they are referred to as mCh-tagged proteins for simplicity.
(E) His$_6$-PomZ ATPase activity. PomY-His$_6$ and DNA was added as indicated. The assay was performed and analysed as in (B).

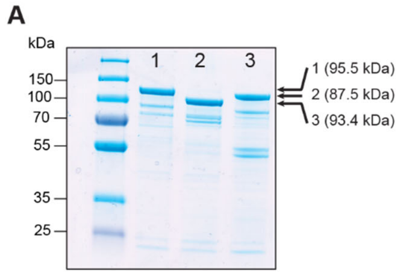


| Number | Protein | Mass [kDa] |
|---|---|---|
| 1 | PomY$^{\Delta 439\text{-}488}$-mCh-His$_6$ | 95.5 |
| 2 | PomY$^{\Delta 496\text{-}623}$-mCh-His$_6$ | 87.5 |
| 3 | PomY$^{\Delta 624\text{-}682}$-mCh-His$_6$ | 93.4 |

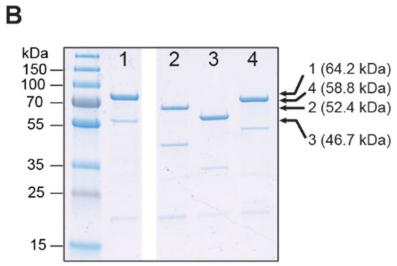


| Number | Protein | Mass [kDa] |
|---|---|---|
| 1 | His$_6$-Sumo-PomY$^{mIDR}$-mCh | 64.2 |
| 2 | His$_6$-Sumo-PomY$^{496\text{-}623}$-mCh | 52.4 |
| 3 | His$_6$-Sumo-PomY$^{624\text{-}682}$-mCh | 46.7 |
| 4 | His$_6$-Sumo-PomY$^{496\text{-}682}$-mCh | 58.8 |

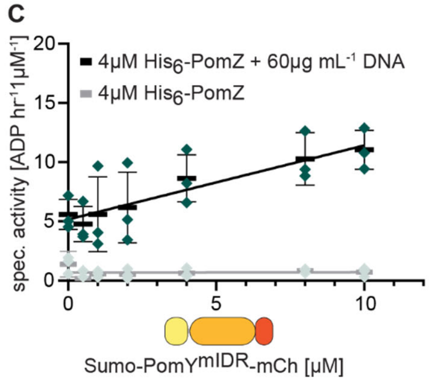


Figure S6. Purification of PomY variants.

(A-B) SDS-PAGE analysis of purified PomY variants. Left lane, molecular size markers with indicated masses. Each protein was assigned a number, which is detailed in the tables below. The proteins in (A) were purified as mCh-His$_6$-tagged proteins; in the main text, they are referred to as mCh-tagged proteins for simplicity. The proteins in (B) were purified as His$_6$-Sumo-tagged proteins; in the main text, they are referred to as Sumo-tagged proteins for simplicity. In B, proteins were separated on the same gel; the gap indicates lanes removed for presentation purposes.

(C) His$_6$-PomZ ATPase activity. Sumo-PomY$^{mIDR}$-mCh and DNA was added as indicated. The assay was performed and analysed as in Fig. 2A.

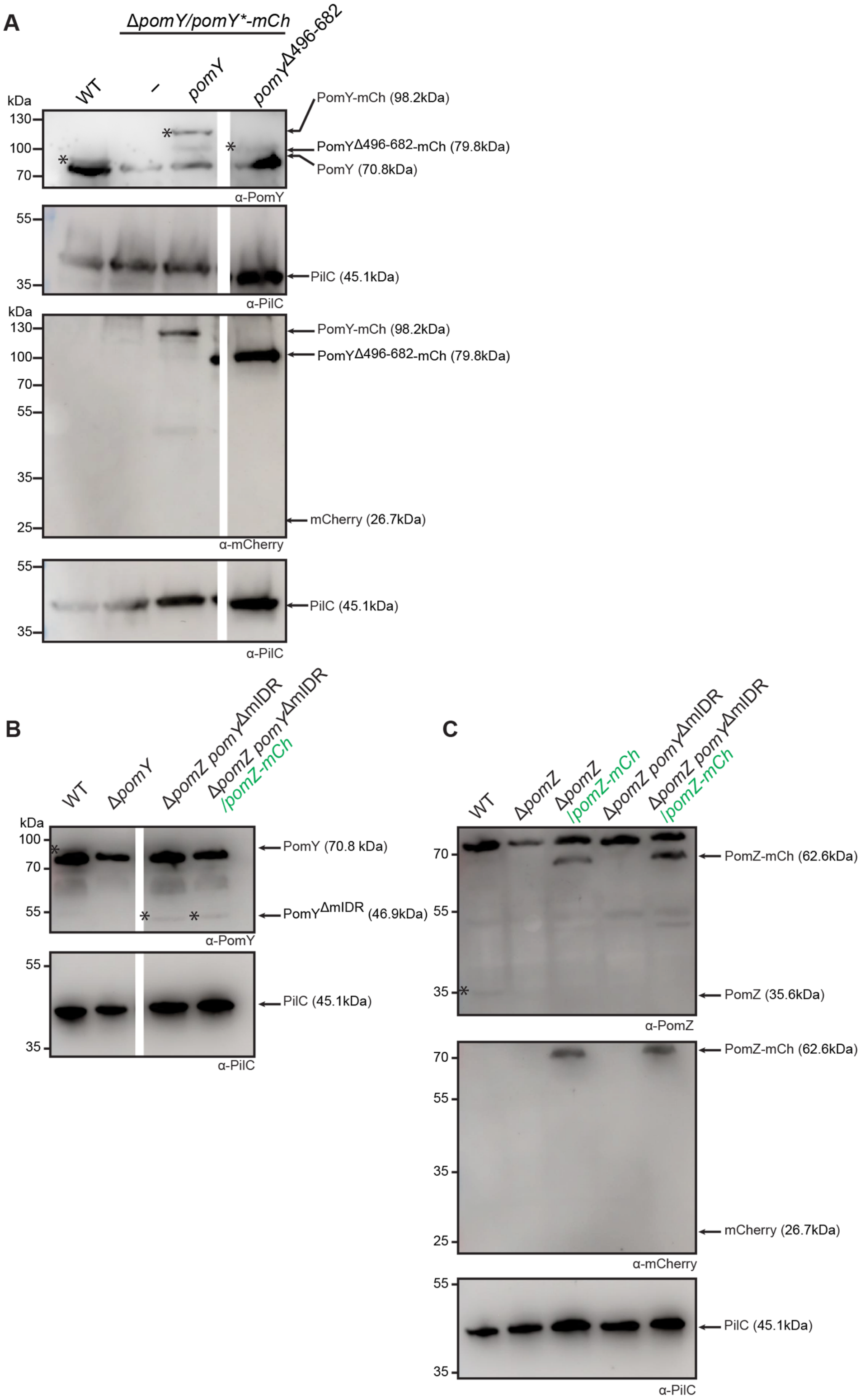


<u>Figure S7. Immunoblot analysis of the accumulation level of PomY variants and PomZ-mCh in *M. xanthus*.</u>

(A) Immunoblot analysis of PomY variants. Total cellular lysates from an equal amount of cells of the indicated *M. xanthus* strains were separated *via* SDS-PAGE, followed by immunoblotting with the indicated antibodies. PilC was used as a loading control. All

samples were analysed on the same blot and the gap indicates lanes that were removed for presentation purposes. *, indicates (from left to right) PomY, PomY-mCh and PomY$^{\Delta 496\text{-}682}$-mCh.
(B) Immunoblot analysis of PomY variants. The analysis was done and presented as in (A). *, indicates (from left to right) PomY, PomY$^{\Delta mIDR}$ and PomY$^{\Delta mIDR}$.
(C) Immunoblot analysis of PomZ-mCh. The analysis was done and presented as in (A). *, indicates PomZ.

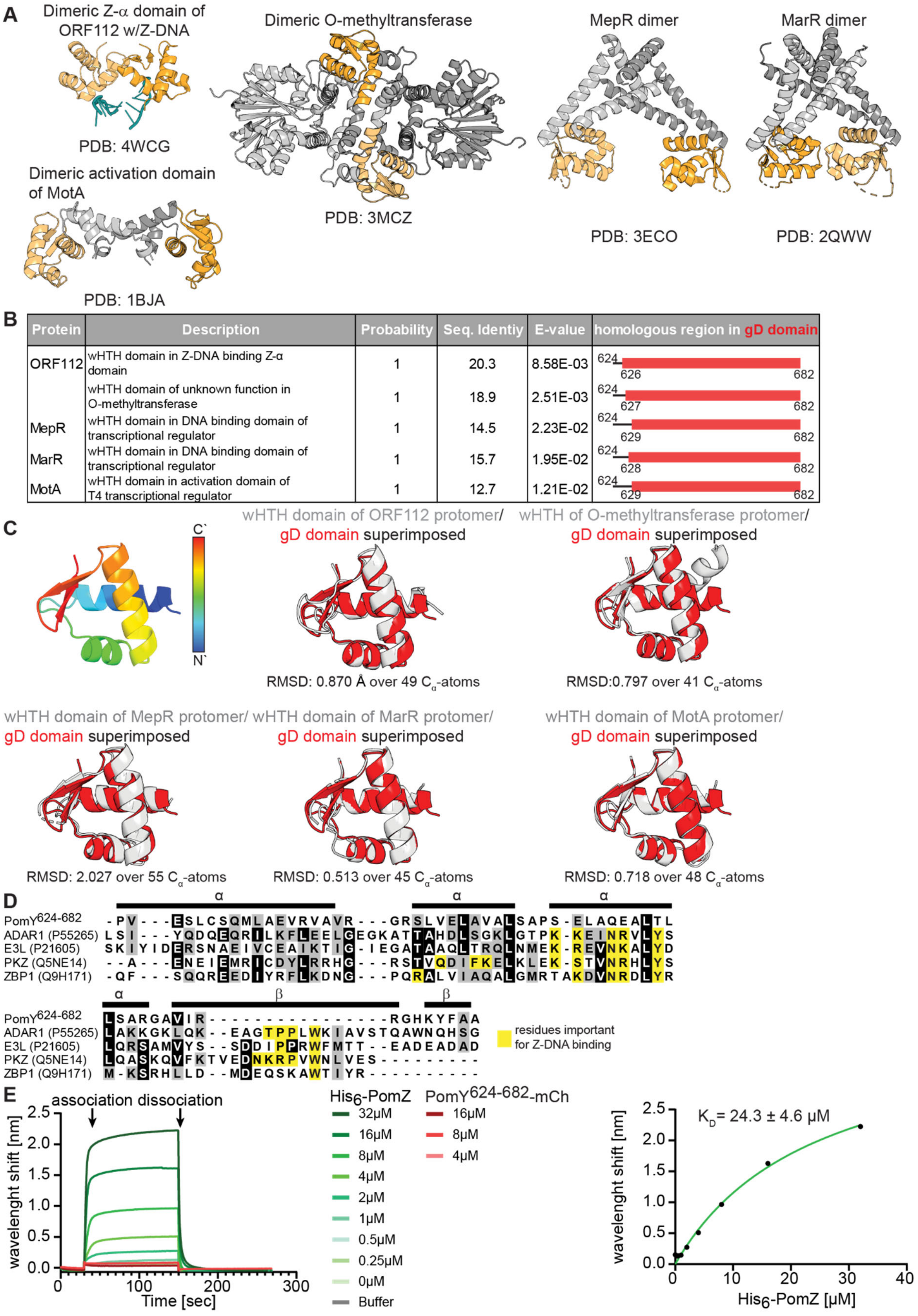


Figure S8. Computational and experimental analysis of the gD domain of PomY.

(A) Solved structures of the five best hits of the Foldseek analysis for structural homologs of the gD domain. The protein regions structurally aligning with the gD domain are marked in orange.
(B) Foldseek analysis of the gD domain against solved structures in the PDB database. From left to right: Protein name of structural homolog, description of the wHTH domain in the structural homologs, probability, sequence identity for structurally aligned residues, E-value; region of the gD domain aligning with the structural homolog.
(C) Structural alignment of the gD domain with the wHTH domains of the five best hits of the FoldSeek analysis. Upper left, the gD domain colored from the N- to the C-terminus; remaining five panels, structural alignment of the gD domain with the wHTH domains n which the gD domain is in red and the structural homologs in light grey.
(D) Multiple sequence alignment of the gD domain with Z-α domains. Secondary structure elements of the gD domain are indicated as black lines above the alignment. Residues involved in binding Z-DNA are marked in yellow (3-6).
(E) BLI analysis of the interaction of the gD domain and $His_6$-PomZ with double stranded DNA. Left panel, a Streptavidin-coated sensor was loaded with a double-biotinylated 226bp DNA fragment and probed with the indicated proteins at the indicated concentrations in the presence of 1mM ATP (association). The protein bound sensor was incubated in wash buffer to observe protein dissociation (dissociation). Note that $PomY^{624\text{-}682}$-mCh only contains the gD domain. Right panel, the wavelength shifts obtained for $His_6$-PomZ at the end of the association phase were plotted as a function of protein concentration and fitted to a one-site specific-binding model. The graph shows the mean of three biological replicates. The $K_D$ value is the mean of the three biological replicates ±STDEV.

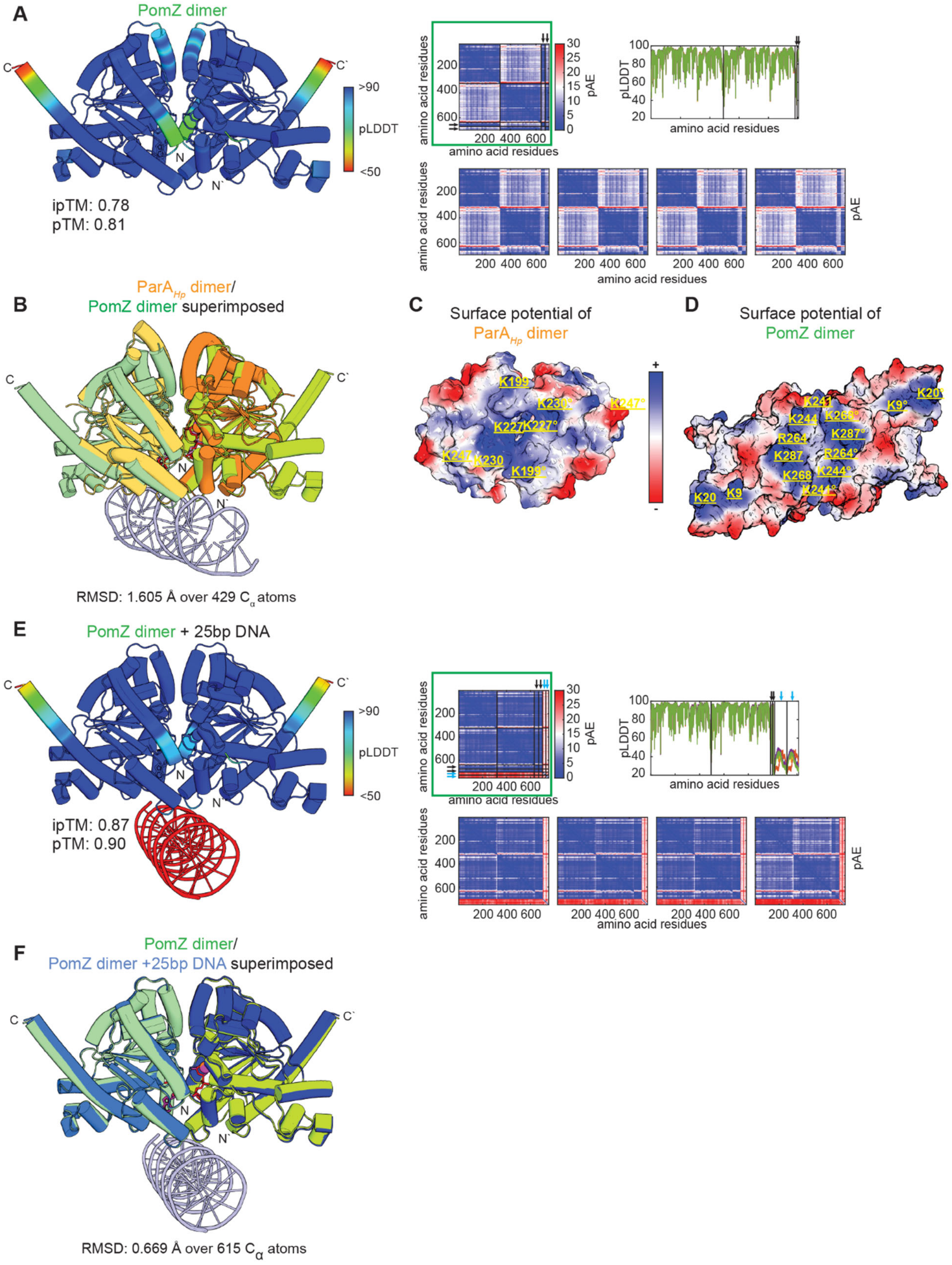


Figure S9. AlphaFold3-based structural model of PomZ dimer.

(A) Left panel, model rank 1 of PomZ dimer colored according to pLDDT score; PomZ was modeled with ATP and $Mg^{2+}$. Right panels, pLDDT plot and pAE plots for the five generated models of the PomZ dimer. The black lines separate the two protomers; black arrows, ATP

and $Mg^{2+}$ molecules. Model rank 1 (highlighted by a green box) was selected for further analyses.
(B) Structural alignment of PomZ dimer with the ATP-bound *Helicobacter pylori* ParA dimer ($ParA_{Hp}$) bound to DNA (7). PomZ dimer is in shades of green, $ParA_{Hp}$ in shades of orange, and DNA in light blue. ATP molecules from PomZ model, red and $Mg^{2+}$, light orange. ADP molecules from $ParA_{HP}$ model, violet and $Mg^{2+}$, magenta.
(C-D) Electrostatic surface potential of $ParA_{Hp}$ dimer (C) and PomZ dimer (D) contoured from +5 to −5 kT $e^{-1}$. Residues marked in yellow are involved in DNA binding in the $ParA_{Hp}$ dimer and suggested to be involved in DNA binding in the PomZ dimer. Residues marked with ° distinguish between the two protomers.
(E) Left panel, model rank 1 of PomZ dimer bound to 25bp DNA fragment colored according to pLDDT values. PomZ was modeled with ATP and $Mg^{2+}$ Right panel, pAE and pLDDT plots of the five models generated. The black lines separate the two protomers; black arrows, ATP and $Mg^{2+}$ molecules, light blue arrows, DNA molecules.
(F) Structural alignment of PomZ dimer modelled in the absence of DNA and PomZ dimer modelled with DNA. PomZ dimer without DNA is in shades of green, PomZ dimer with DNA in blue. DNA is coloured in pale blue. ATP molecules from the PomZ model without DNA, red and $Mg^{2+}$ ions, light orange; ATP molecules of the PomZ model with DNA, purple and $Mg^{2+}$ ions, magenta.

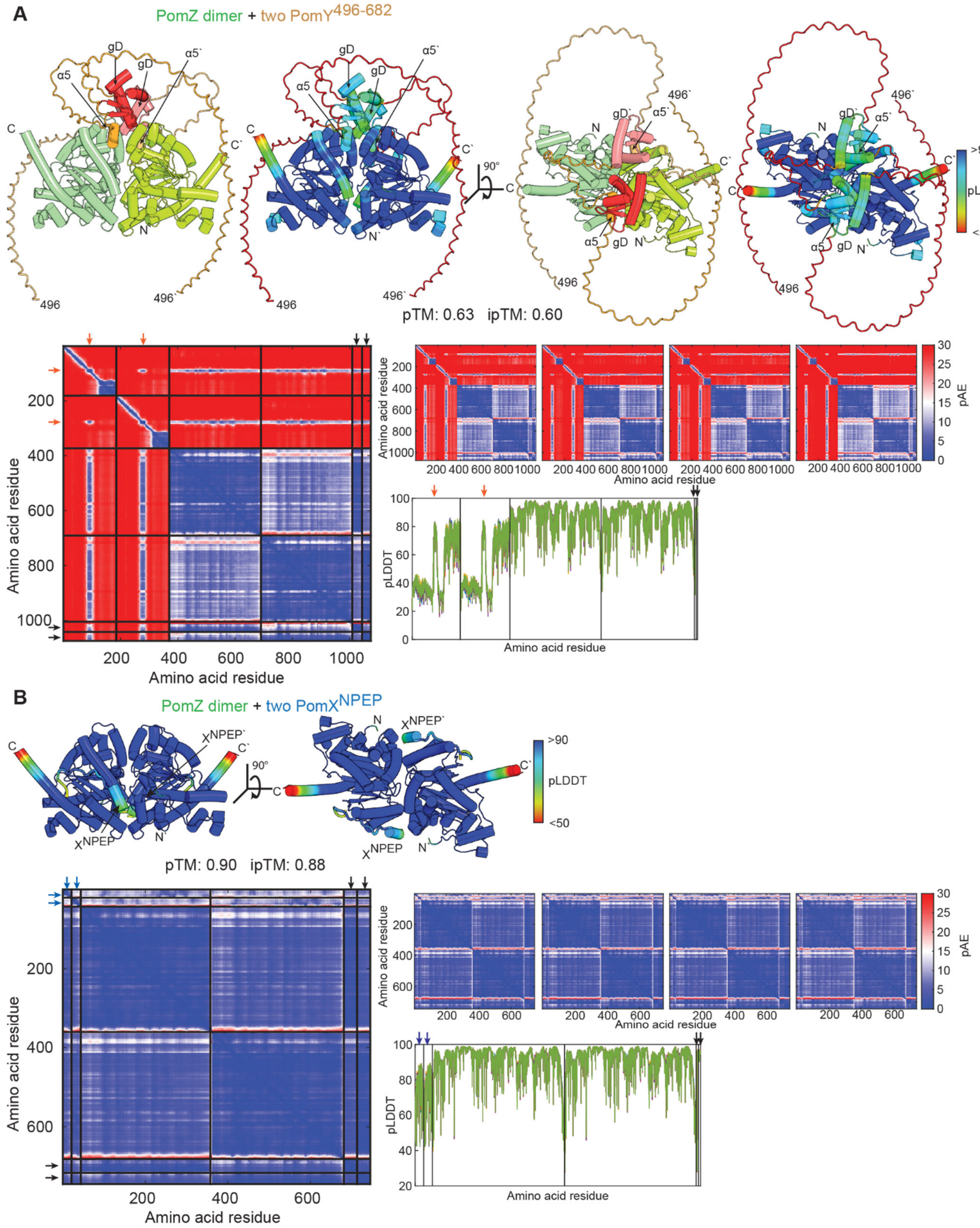


<u>Figure S10. AlphaFold3-based structural model of PomZ dimer in complex with PomY$^{496\text{-}682}$ or PomX$^{NPEP}$.</u>

(A) Top panels, model rank 1 of PomZ dimer with two molecules of PomY$^{496\text{-}682}$. In the left panels, the model is shown with the PomZ dimer in shades of green, the IDRs in PomY$^{496\text{-}682}$ in brown, α5 in orange and the gD domain in red. In the right panels, the model is colored

according to pLDDT score. PomZ was modeled with ATP and $Mg^{2+}$ but the ATP molecules and $Mg^{2+}$ are not included for simplicity. Lower panels, pLDDT plot and pAE plots for the five generated models. The black lines separate the two PomZ protomers and the two $PomY^{496-682}$ molecules; black arrows, ATP and $Mg^{2+}$, and orange arrows α5 of $PomY^{496-682}$. Model rank 1 is shown in high magnification and was selected for further analyses. Note that the model is of low confidence based on pTM and ipTM scores. The large proportion of IDRs in $PomY^{496-682}$, which have low pLDDT scores, causes these low values.
(B) Top panels, model rank 1 of PomZ dimer with two molecules of $PomX^{NPEP}$ colored according to pLDDT score. PomZ was modeled with ATP and $Mg^{2+}$ but the ATP molecules and $Mg^{2+}$ are not included for simplicity. Lower panels, pLDDT plot and pAE plots for the five generated models. The black lines separate the two PomZ protomers and two $PomX^{NPEP}$ molecules; black arrows, ATP and $Mg^{2+}$, and blue arrows $PomX^{NPEP}$. Model rank 1 is shown in high magnification and was selected for further analyses.

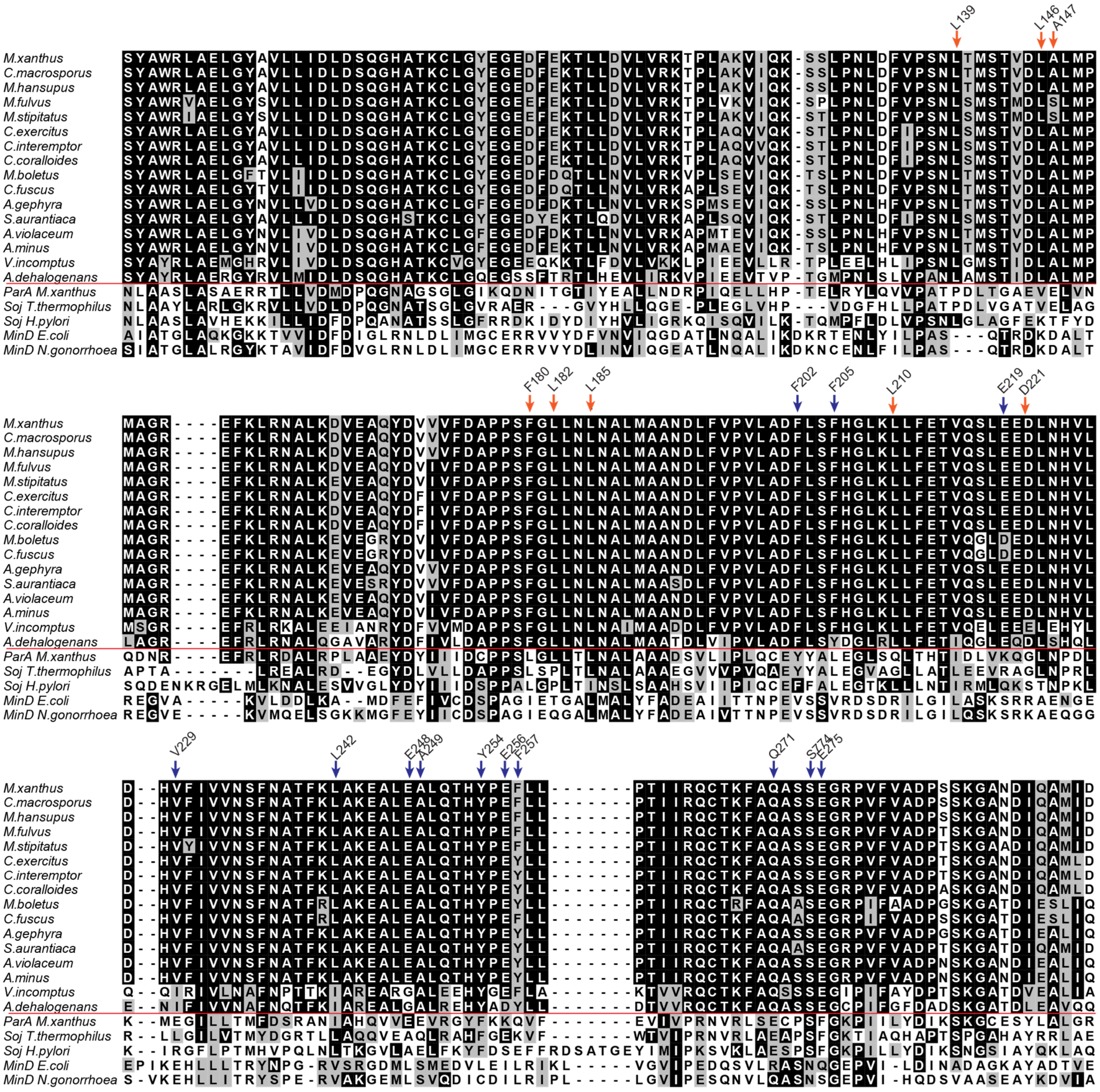


Figure S11. Multiple sequence alignment of PomZ orthologs from Myxobacteria with fully sequenced genomes as well as ParA orthologs and MinD orthologs.

The alignment is visualized with a shading threshold of 80% based on the BLOSUM62 matrix. Grey boxes indicate similarity and black boxes identity. Orange arrows indicate residues in PomZ of *M. xanthus* predicted to interact with α5 of PomY$^{496-682}$. Blue arrows indicate residues predicted to interact with PomX$^{NPEP}$.

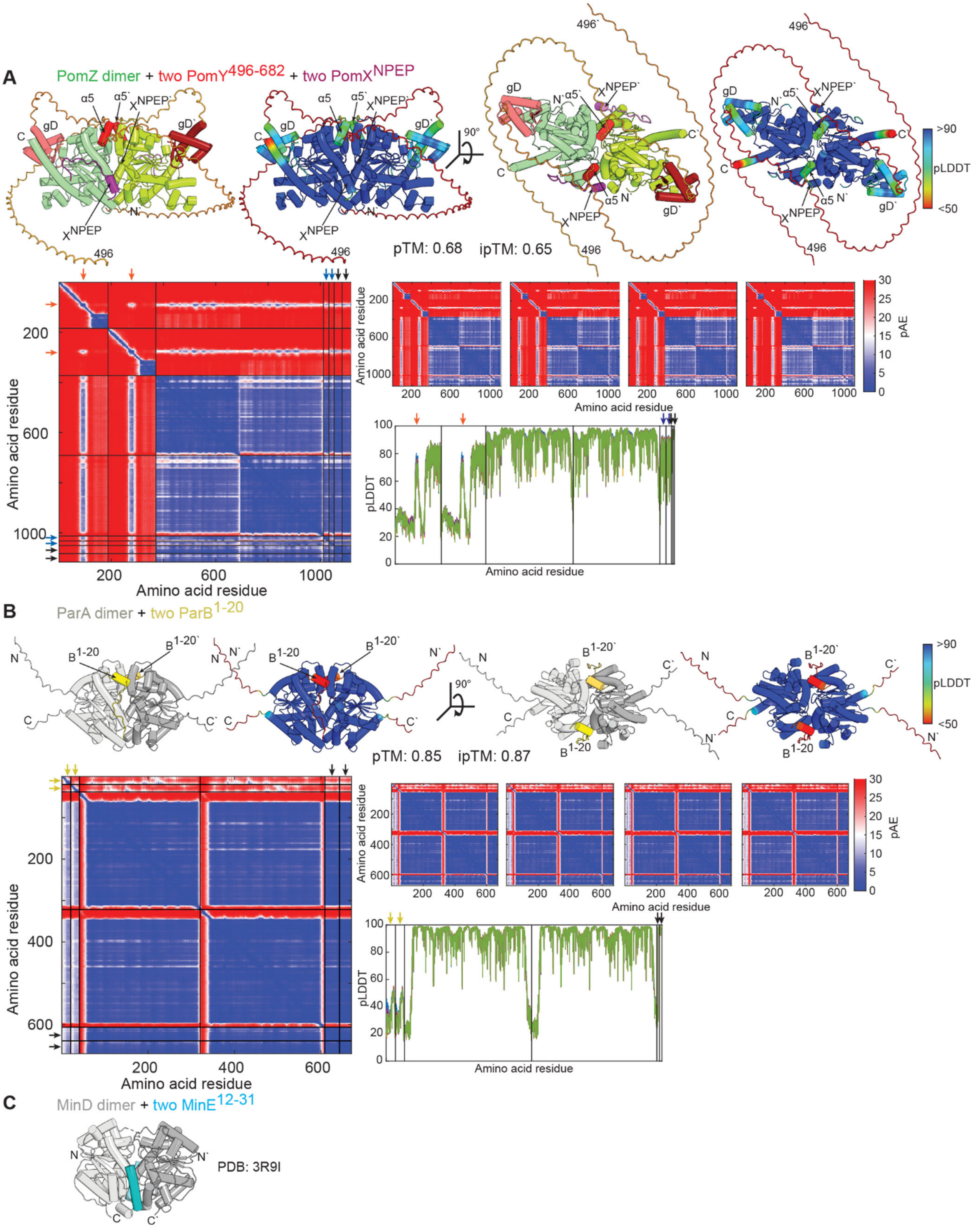


Figure S12. AlphaFold3-based structural model of the PomZ/PomY$^{496-682}$/PomX$^{NPEP}$ interactions

(A) Top panels, model rank 1 of PomZ dimer with two molecules of $PomY^{496-682}$ and two molecules of $PomX^{NPEP}$. In the left panels, the model is shown with the PomZ dimer in shades of green, the IDRs in $PomY^{496-682}$ in brown, α5 in red, the gD domain in dark red, and $PomX^{NPEP}$ in purple. In the right panels, the model is colored according to pLDDT score. PomZ was modeled with ATP and $Mg^{2+}$ but the ATP molecules and $Mg^{2+}$ are not included for simplicity. Lower panels, pLDDT plot and pAE plots for the five models generated. The black lines separate the two PomZ protomers, the two $PomY^{496-682}$ molecules and the two $PomX^{NPEP}$ molecules; black arrows, ATP and $Mg^{2+}$, red arrows α5 of $PomY^{496-682}$, and purple arrows $PomX^{NPEP}$. Model rank 1 is shown in high magnification and was selected for further analyses. Note that the model is of low confidence based on pTM and ipTM scores. The large proportion of IDRs in $PomY^{496-682}$, which have low pLDDT scores, causes these low values.
(B) Top panels, model rank 1 of ParA dimer with two molecules $ParB^{1-20}$. In the left panels, the model is shown with the ParA dimer in shades of grey, and the two $ParB^{1-20}$ molecules in yellow In the right panels, the model is colored according to pLDDT score. ParA was modeled with ATP and $Mg^{2+}$ but the ATP molecules and $Mg^{2+}$ are not included for simplicity. Lower panels, pLDDT plot and pAE plots for the five generated models. The black lines separate the two ParA protomers and the two $ParB^{1-20}$ molecules; black arrows, ATP and $Mg^{2+}$, and yellow arrows $ParB^{1-20}$. Model rank 1 is shown in high magnification and was selected for further analyses. The ParA and ParB proteins analyzed are those of *M. xanthus* and are involved in chromosome segregation (8).
(C) Solved structure of the *E. coli* MinD dimer (grey shades) with two molecules of $MinE^{12-31}$ (cyan) (9).

**Supplementary Table 1.** *M. xanthus* strains used in this work.

| Strain | Relevant charateristics [1] | Reference |
|---|---|---|
| SA4420 | Δ*mglA* | (10) |
| SA4779 | Δ*mglA* Δ*pomY* | (1) |
| SA7064 | Δ*mglA* Δ*pomY*/*attB*::$P_{nat}$ *pomY-mCh* (pDS8) | (1) |
| SA11833 | Δ*mglA* Δ*pomY*/*attB*::$P_{nat}$ *pomY*$^{\Delta 496\text{-}682}$-*mCh* (pPK82) | This study |
| SA3108 | Δ*pomZ* | (11) |
| SA4772 | Δ*mglA* Δ*pomZ* | This study |
| SA4798 | Δ*mglA* Δ*pomZ*/*attB*::$P_{nat}$ *pomZ-mCh* (pKA28) | This study |
| SA11827 | Δ*mglA* Δ*pomZ* *pomY*$^{\Delta mIDR}$ | This study |
| SA11828 | Δ*mglA* Δ*pomZ* *pomY*$^{\Delta mIDR}$/*attB*::$P_{nat}$ *pomZ-mCh* (pKA28) | This study |

[1] $P_{nat}$ indicates that genes were expressed from their native promoter.

[2] Plasmids integrated in a single copy at the Mx8 *attB* site are listed in brackets.

**Supplementary Table 2.** Plasmids used in this work

| Plasmid | Relevant characteristics [1] | Reference |
|---|---|---|
| pET24b+ | Expression vector for $His_6$-tagged proteins | Novagen |
| pTB146 | $P_{T7}$-$His_6$-Sumo; $Amp^R$ | (12) |
| pBJ114 | $Km^R$; *galK*$^+$; vector for generating in-frame deletions; $Km^R$ | (13) |
| pSL16 | Δ*mglA*; *galK*$^+$; $Km^R$ (for in-frame deletion of *mglA*) | (10) |
| pSWU30 | *attP*; $Tet^R$ | (14) |
| pSW105 | $P_{pilA}$; *attP*; $Km^R$ | (15) |
| pMR3690 | Vector for pAH204 | (16) |
| pKNT25 | Vector for pDS52 and pDS64 | (17) |
| pKA3 | Overexpression of $His_6$-PomZ; $Amp^R$ | (11) |
| pDS3 | Overexpression of PomY-$His_6$; $Km^R$ | (2) |
| pAH194 | Overexpression of PomY-mCh-$His_6$; $Km^R$ | (1) |
| pAH196 | Overexpression of $PomY^{CC}$-mCh-$His_6$; $Km^R$ | (1) |
| pAH197 | Overexpression of $PomY^{HEAT}$-mCh-$His_6$; $Km^R$ | (1) |
| pAH198 | Overexpression of $PomY^{mIDR}$-mCh-$His_6$; $Km^R$ | (1) |
| pAH187 | Overexpression of $PomY^{CC\_HEAT}$-mCh-$His_6$; $Km^R$ | (1) |
| pAH195 | Overexpression of $PomY^{HEAT\_mIDR}$-mCh-$His_6$; $Km^R$ | (1) |
| pAH217 | Overexpression of $PomY^{CC\_mIDR}$-mCh-$His_6$; $Km^R$ | This study |
| pPK57 | Overexpression of $PomY^{\Delta 438\text{-}488}$-mCh-$His_6$; $Km^R$ | This study |
| pPK67 | Overexpression of $PomY^{\Delta 496\text{-}623}$-mCh-$His_6$; $Km^R$ | This study |
| pPK68 | Overexpression of $PomY^{\Delta 624\text{-}682}$-mCh-$His_6$; $Km^R$ | This study |
| pPK83 | Overexpression of $His_6$-Sumo-$PomY^{mIDR}$-mCh; $Amp^R$ | This study |
| pPK84 | Overexpression of $His_6$-Sumo-$PomY^{496\text{-}623}$-mCh; $Amp^R$ | This study |

| | | |
|---|---|---|
| pPK85 | Overexpression of $His_6$-Sumo-PomY$^{624\text{-}682}$-mCh; $Amp^R$ | This study |
| pPK86 | Overexpression of $His_6$-Sumo-PomY$^{496\text{-}682}$-mCh; $Amp^R$ | This study |
| pPK72 | Overexpression of PomY$^{624\text{-}682}$-mCh-$His_6$; $Km^R$ | This study |
| pGBT9 | Y2H plasmid expressing Gal4BD | Clontech |
| pDS82 | Y2H plasmid expressing Gal4BD-PomY | (2) |
| pDS139 | Y2H plasmid expressing Gal4BD-PomY$^{CC_HEAT}$ | This study |
| pAH274 | Y2H plasmid expressing Gal4BD-PomY$^{mIDR}$ | This study |
| pGAD424 | Y2H plasmid expressing Gal4AD | Clontech |
| pDS83 | Y2H plasmid expressing Gal4AD-PomZ | (2) |
| pDS52 | PomZ$^{G62V}$; $Km^R$<br>(template for pPK90) | This study |
| pDS64 | PomZ$^{K268E}$; $Km^R$<br>(template for pPK91) | This study |
| pKA12 | $His_6$-PomZ$^{D90A}$; $Amp^R$<br>(template for pPK89) | This study |
| pPK90 | Y2H plasmid expressing Gal4AD-PomZ$^{G62V}$ | This study |
| pPK91 | Y2H plasmid expressing Gal4AD-PomZ$^{K268E}$ | This study |
| pPK89 | Y2H plasmid expressing Gal4AD-PomZ$^{D90A}$ | This study |
| pDS8 | $P_{nat}$ PomY-mCh; *attP*; $Tet^R$ | (2) |
| pPK82 | $P_{nat}$ PomY$^{\Delta 496\text{-}682}$-mCh; *attP*; $Tet^R$ | This study |
| pPK78 | PomY$^{\Delta mIDR}$; *galK*$^+$; $Km^R$<br>(for in-frame deletion of mIDR) | This study |
| pKA28 | $P_{nat}$ PomZ-mCh; *attP*; $Tet^R$ | (11) |
| pDS7 | $P_{pilA}$-PomY-mCh, $Km^R$<br>(template for pPK35 & pPK53) | (2) |
| pAH204 | $P_{pilA}$ PomY$^{CC_mIDR}$-mCh; *attP*; $Km^R$<br>(template for pAH217) | This study |
| pPK53 | $P_{pilA}$ PomY$^{\Delta 438\text{-}488}$-mCh; *attP*; $Km^R$<br>(template for pPK57) | This study |

| | | |
|---|---|---|
| pPK61 | $P_{pilA}$ PomY$^{mIDR}$-mCh; *attP*; Km$^R$ (template for pPK67) | This study |
| pPK38 | $P_{pilA}$ PomY$^{\Delta 624-682}$-mCh; *attP*; Km$^R$ (template for pPK68) | This study |
| pPK71 | PomY$^{496-623}$-mCh-His$_6$; Km$^R$ (template for pPK84) | This study |
| pPK74 | PomY$^{496-682}$-mCh-His$_6$; Km$^R$ (template for pPK86) | This study |
| pPK35 | $P_{pilA}$-PomY$^{CC-mIDR}$-mCh; *attP*; Km$^R$ (template for pAH217) | This study |
| pAH107 | PomZ$^{G62V}$; Km$^R$ (template for pPK90) | This study |
| pAH110 | PomZK268E ; KmR (template for pPK91) | This study |
| pDS43 | PomZ-mCh; Km$^R$ (template for pAH107 & pAH110) | (2) |

[1] $P_{nat}$ and $P_{pilA}$ indicate the native promoter and the *pilA* promoter, respectively.

**Supplementary Table 3.** Oligonucleotides used in this work

| Oligo-nucleotide | Sequence (5'-3') |
|---|---|
| DS14 | GCGCATATGAGCGACGAGCGTCCG |
| DS274 | GCGAAGCTTCTTGTACAGCTCGTCCATGC |
| mcherry stop rev | GGGAAGCTTTTACTTGTACAGCTCGTC |
| PK87 | CGATATTATTGAGGCTCACAGAGAACAGATTGGTGGTATGCTGGAGCAGCCTCGTCCGGTGGTGGCTTCGG |
| PK86 | CCTTTCGGGCTTTGTTAGCAGCCGGATCCTCATCACTTGTACAGCTCGTCCATGCCGCCGGTGGAGTGG |
| PK85 | CGATATTATTGAGGCTCACAGAGAACAGATTGGTGGTATGCCGGTGGGGCCTGTGCAGGGGCGAGC |
| PK88 | CGATATTATTGAGGCTCACAGAGAACAGATTGGTGGTATGGCGTCACCGGTGGAATCCCTCTGCTCGCAGATGC |
| PK85 | CGATATTATTGAGGCTCACAGAGAACAGATTGGTGGTATGCCGGTGGGGCCTGTGCAGGGGCGAGCAGGCCCC |
| AH230 | GCGGAATTCCTGGAGCAGCCTCGTCCGG |
| AH231 | GCGAGATCTTCAAGCGGCGAAGTATTTGTG |
| Mxan0635 fwd SalI linker | GCGGTCGACCTATGGAAGCGCCGACGTACAGC |
| Mxan0635 rev stop BglII | GCGAGATCTTCAGCCGGCCTGCTGGGTGCC |
| DS1 | CCGGAATTCCGACGAGCAGTTGAGCACCAG |
| PK84 | CCGGCGGCGGATCTGGCGGAACCAGCGCCCTGCGCGAGGC |
| PK83 | GCCTCGCGCAGGGCGCTGGTTCCGCCAGATCCGCCGCCGG |
| mcherry stop rev HindIII | GGGAAGCTTTTACTTGTACAGCTCGTC |
| PK64 | CGCAAGCTTCGCTGCTCCAGGCCTTCGC |
| PK65 | CGGCTACCTTGCCTCACCGCGCCCGCACAGGC |
| PK66 | GCCTGTGCGGGCGCGGTGAGGCAAGGTAGCCG |
| PK80 | GCGGAATTCCCGCTGCACCTCC |
| PK77 | GCGCATATGGCGTCACCGGTGGAATCCC |
| Mxan0634 fwd EcoRI | GCGGAATTCGTGAGCGACGAGCGTCCGGAC |
| Mxan0634 HEAT rev BglII | GCGAGATCTTCACCGCGCCCGCACAGGC |
| cluster3 prom 1 bioteg | GGGCCACCAAGCCCACCGCC |
| cluster3 prom 2 bioteg | GGCAAATTCCCAGGAGGAGG |
| DS40 | GCGGAATTCTTACTTGTACAGCTCGTC |
| DS11 | CAGCGCTTCCTGCAGGCGCTGGAGCAGCCTCGTCCG |
| DS4 | GCGTCTAGAGTGAGCGACGAGCGTCCG |
| PK39 | CTGGAGCAGCCTCGTGGCCTCGCGCAGGGC |
| PK40 | GCCCTGCGCGAGGCCACGAGGCTGCTCCAG |
| PK44 | GCGCAGGGCGCTGGTGTGGAATCCCTCTGC |

| | |
|---|---|
| PK45 | GCAGAGGGATTCCACACCAGCGCCCTGCGC |
| PK78 | GCGCATATGCCGGTGGGGCCTGTGCAGGG |
| PK21 | CGCAGATCTGGCGGATGACGCACTGGGGCGCGAGGGGGC |
| KA419 | CTCATCGACCTCGCCAGCCAGGGCCAC |
| KA420 | GTGGCCCTGGCTGGCGAGGTCGATGAG |
| Mxan_0635-4 pomZ fwd | GCGTCTAGAGATGGAAGCGCCGACGTAC |
| Mxan_0635-5 pomZ rev | GCGGGTACCCGGCCGGCCTGCTGGGTGCC |
| KA413 | TCCTGAACTTCAAGGTTGGCACCGGCAAGAC |
| KA414 | GTCTTGCCGGTGCCAACCTTGAAGTTCAGGA |
| AH91 | ATCCGGCAGTGCACCGAGTTCGCGCAGGCCTCC |
| AH92 | GGAGGCCTGCGCGAACTCGGTGCACTGCCGGAT |

**Supplementary Table 4.** Construction of plasmids

| Plasmid | Construction |
|---|---|
| pAH217 | A fragment containing PomY$^{CC}$ was amplified from pDS7, which contains PomY-mCh, with the primers DS4 and DS10. A second fragment containing PomY$^{mIDR}$-mCh was amplified from pDS7 with the primers DS11 and mCh stop rev HindIII. The two fragments were fused with the primers DS4 and mCh stop rev HindIII in a PCR. The fragment and vector pSW105 were digested with XbaI and HindIII and, after dephosphorylation, used for ligation. Competent NEB Turbo cells were transformed with the ligation reaction, and the obtained plasmid pPK35 was sequenced. Subsequently, a fragment containing PomY$^{CC\text{-}mIDR}$-mCh was amplified from pPK35, which contains PomY$^{CC\text{-}mIDR}$-mCh, with the Primers DS14 and DS40. The fragment and vector pMR3690 were digested with NdeI and EcoRI and, after dephosphorylation, used for ligation. Competent NEB Turbo cells were transformed with the ligation reaction, and the obtained plasmid pAH204 was sequenced. Subsequently, the fragment containing PomY$^{CC\text{-}mIDR}$-mCh was amplified from pAH204 with the primers DS14 and DS274. The fragment and vector pET24b+ were digested with NdeI and HindIII and, after dephosphorylation, used for ligation. Competent NEB Turbo cells were transformed with the ligation reaction, and the obtained plasmid pAH217 was sequenced. |
| pPK57 | A fragment containing PomY$^{1\text{-}437}$ was amplified from pDS7, which contains PomY-mCh, with the primers DS4 and PK40. A second fragment containing PomY$^{489\text{-}682}$-mCh was amplified from pDS7 with the primers PK39 and mCh stop rev HindIII. The two fragments were fused with the primers DS4 and mCh stop rev HindIII in a PCR. The fragment and vector pSW105 were digested with XbaI and HindIII and, after dephosphorylation, used for ligation. Competent NEB Turbo cells were transformed with the ligation reaction, and the obtained plasmid pPK53 was sequenced. Subsequently, the fragment containing PomY$^{\Delta 438\text{-}488}$-mCh was amplified from pPK53 with the primers DS14 and mCh stop rev. The fragment and vector pET24b+ were digested with NdeI and HindIII and, after dephosphorylation, used for ligation. Competent NEB Turbo cells were transformed with the ligation reaction, and the obtained plasmid pPK57 was sequenced. |
| pPK67 | A fragment containing PomY$^{1\text{-}495}$ was amplified from pDS8, which contains PomY-mCh, with the primers DS4 and PK45. A second fragment containing PomY$^{624\text{-}682}$-mCh was amplified from pD8 with the primers PK44 and mCh stop rev HindIII. The two fragments were fused with the primers DS4 and mCh stop rev HindIII in a PCR. The fragment and vector pSW105 were digested with XbaI and HindIII and, after dephosphorylation, used for ligation. Competent NEB Turbo cells were transformed with the ligation reaction, and the obtained plasmid pPK61 was sequenced. Subsequently, the fragment containing PomY$^{\Delta 496\text{-}623}$-mCh was amplified from pPK61 with the primers DS14 and DS274. The fragment and vector pET24b+ were digested with NdeI and HindIII and, after dephosphorylation, used for ligation. Competent NEB Turbo cells were transformed with the ligation reaction, and the obtained plasmid pPK67 was sequenced. |
| pPK68 | A fragment containing PomY$^{\Delta 624\text{-}682}$ was amplified from pDS8, which contains PomY-mCh, with the Primers DS4 and PK21. The fragment was digested with XbaI and BglII, and the vector pSWU30 was digested with XbaI and BamHI and, after dephosphorylation, used for ligation. Competent NEB Turbo cells were transformed with the ligation reaction, and the obtained plasmid pPK38 was sequenced. Subsequently, the fragment containing PomY$^{\Delta 624\text{-}682}$-mCh was amplified from pPK38 with the primers DS14 and DS274. The fragment and vector pET24b+ were digested with NdeI and HindIII and, after dephosphorylation, used for ligation. Competent NEB Turbo cells were transformed with the ligation reaction, and the obtained plasmid pPK68 was sequenced. |
| pPK83 | A fragment containing PomY$^{mIDR}$-mCh was amplified from pAH198, which contains PomY$^{mIDR}$-mCh, with the Primers PK87 and PK86. The Fragment was |

| | |
|---|---|
| | inserted into the vector pTB146 with the Gibson Assembly Cloning Kit, NEB #E5510S. |
| pPK84 | A fragment containing PomY$^{496-623}$-mCh was amplified from pPK68, which contains PomY$^{\Delta 624-682}$-mCh, with the Primers PK78 and DS274. The fragment and vector pET24b+ were digested with NdeI and HindIII and, after dephosphorylation, used for ligation. Competent NEB Turbo cells were transformed with the ligation reaction, and the obtained plasmid pPK71 was sequenced. Subsequently, the fragment containing PomY$^{496-682}$-mCh was amplified from pPK71 with the primers PK85 and PK86. The Fragment was inserted into the vector pTB146 with the Gibson Assembly Cloning Kit, NEB #E5510S. |
| pPK85 | A fragment containing PomY$^{624-682}$-mCh was amplified from pAH194, which contains His$_6$-PomY-mCh, with the Primers PK77 and DS274. The fragment and vector pET24b+ were digested with NdeI and HindIII and, after dephosphorylation, used for ligation. Competent NEB Turbo cells were transformed with the ligation reaction, and the obtained plasmid pPK72 was sequenced. Subsequently, the fragment containing PomY$^{624-682}$-mCh was amplified from pPK72 with the primers PK88 and PK86. The Fragment was inserted into the vector pTB146 with the Gibson Assembly Cloning Kit, NEB #E5510S. |
| pPK86 | A fragment containing PomY$^{496-682}$-mCh was amplified from pAH194, which contains His$_6$-PomY-mCh, with the Primers PK78 and DS274. The fragment and vector pET24b+ were digested with NdeI and HindIII and, after dephosphorylation, used for ligation. Competent NEB Turbo cells were transformed with the ligation reaction, and the obtained plasmid pPK74 was sequenced. Subsequently, the fragment containing PomY$^{496-682}$-mCh was amplified from pPK74 with the primers PK85 and PK86. The Fragment was inserted into the vector pTB146 with the Gibson Assembly Cloning Kit, NEB #E5510S. |
| pDS139 | A fragment containing PomY$^{CC-HEAT}$ was amplified from DK1622 genomic DNA with the Primers Mxan 0634 fwd EcoRI and Mxan0634 HEAT rev BglII. The fragment was digested with EcoRI and BglII, and the vector pGBT9 was digested with XbaI and BamHI and, after dephosphorylation, used for ligation. Competent NEB Turbo cells were transformed with the ligation reaction, and the obtained plasmid pDS139 was sequenced. |
| pAH274 | A fragment containing PomY$^{mIDR}$ was amplified from DK1622 genomic DNA with the Primers AH230 and AH231. The fragment and vector pGBT9 were digested with EcoRI and BamHI and, after dephosphorylation, used for ligation. Competent NEB Turbo cells were transformed with the ligation reaction, and the obtained plasmid pAH274 was sequenced. |
| pPK90 | The G62V point mutation in PomZ was inserted by amplifying the plasmid pDS43 with the primers KA413 and KA414, carrying the point mutation, with the QuikChange II XL Site-Directed Mutagenesis Kit (Agilent), and the obtained plasmid pAH107 was sequenced. Subsequently, a fragment containing PomZ$^{G62V}$ was amplified from pAH107, which contains PomZ$^{G62V}$, with the Primers Mxan_0635-4 pomZ fwd and Mxan_0635-5 pomZ rev. The fragment and vector pKNT25 were digested with XbaI and KpnI and, after dephosphorylation, used for ligation. Competent NEB Turbo cells were transformed with the ligation reaction, and the obtained plasmid pDS52 was sequenced. Subsequently, the fragment containing PomZ$^{G62V}$ was amplified from pDS52 with the primers Mxan 0634 fwd SalI linker and Mxan0635 rev stop BglII. The fragment and vector pGAD424 were digested with SalI and BglII and, after dephosphorylation, used for ligation. Competent NEB Turbo cells were transformed with the ligation reaction, and the obtained plasmid pPK90 was sequenced. |
| pPK91 | The K268E point mutation in PomZ was inserted by amplifying the whole plasmid pDS43 with the primers AH91 and AH92, carrying the point mutation, with the QuikChange II XL Site-Directed Mutagenesis Kit (Agilent), and the obtained plasmid pAH110 was sequenced. Subsequently, a fragment containing PomZ$^{K268E}$ was amplified from pAH110, which contains PomZ$^{K268E}$, with the |

| | |
|---|---|
| | Primers Mxan_0635-4 pomZ fwd and Mxan_0635-5 pomZ rev. The fragment and vector pKNT25 were digested with XbaI and KpnI and, after dephosphorylation, used for ligation. Competent NEB Turbo cells were transformed with the ligation reaction, and the obtained plasmid pDS64 was sequenced. Subsequently, the fragment containing PomZ$^{K268E}$ was amplified from pDS64 with the primers Mxan 0634 fwd SalI linker and Mxan0635 rev stop BglII. The fragment and vector pGAD424 were digested with SalI and BglII and, after dephosphorylation, used for ligation. Competent NEB Turbo cells were transformed with the ligation reaction, and the obtained plasmid pPK91 was sequenced. |
| pPK89 | The D90A point mutation in PomZ was inserted by amplifying the whole plasmid pKA28 with the primers KA419 and KA420, carrying the point mutation, with the QuikChange II XL Site-Directed Mutagenesis Kit (Agilent), and the obtained plasmid pKA12 was sequenced. Subsequently, the fragment containing PomZ$^{D90A}$ was amplified from pDS64 with the primers Mxan 0634 fwd SalI linker and Mxan0635 rev stop BglII. The fragment and vector pGAD424 were digested with SalI and BglII and, after dephosphorylation, used for ligation. Competent NEB Turbo cells were transformed with the ligation reaction, and the obtained plasmid pPK89 was sequenced. |
| pPK82 | A fragment containing PomY$^{1\text{-}495}$ was amplified from pDS8, which contains PomY-mCh, with the primers DS1 and PK84. A second fragment containing mCh was amplified from pD8 with the primers PK83 and mCh stop rev HindIII. The two fragments were fused with the primers DS1 and mCh stop rev HindIII in a PCR. The fragment and vector pSWU30 were digested with EcoRI and HindIII and, after dephosphorylation, used for ligation. Competent NEB Turbo cells were transformed with the ligation reaction, and the obtained plasmid pPK82 was sequenced. |
| pPK78 | A fragment containing 500bp upstream of PomYmIDR was amplified from pAH194, which contains PomY-mCh, with the primers PK64 and PK65. A second fragment containing 500bp of the genomic region immediately following the *pomY* locus, was amplified from genomic SA4772 DNA with the primers PK66 and PK80. The two fragments were fused with the primers PK64 and PK80 in a PCR. The fragment and vector pBJ114 were digested with EcoRI and HindIII and, after dephosphorylation, used for ligation. Competent NEB Turbo cells were transformed with the ligation reaction, and the obtained plasmid pPK78 was sequenced. |
| pPK72 | A fragment containing PomY$^{624\text{-}682}$-mCh was amplified from pAH194, which contains PomY-mCh-His$_6$, with the Primers PK77 and DS274. The fragment and vector pET24b+ were digested with NdeI and HindIII and, after dephosphorylation, used for ligation. Competent NEB Turbo cells were transformed with the ligation reaction, and the obtained plasmid pPK72 was sequenced. |
